# Skull and scalp segmentation in neonatal cerebral MRI using subject-specific probability models

**DOI:** 10.1101/2022.05.06.490211

**Authors:** Elham Hokmabadi, Hamid Abrishami Moghaddam, Mehrana Mohtasebi, Amirreza Kazemloo, Masume Gity, Fabrice Wallois

## Abstract

This study presents a new approach for segmenting cranial bones in magnetic resonance images (MRIs) acquired from neonates in the gestational age range of 39 to 42 weeks. the approach uses subject-specific probability maps of the skull and scalp, created from atlas computed tomography (CT) images taken retrospectively from neonates in the same age range. the method also uses a subject-specific probability map of cerebrospinal fluid (CSF), constructed from retrospective atlas MRIs. To build skull, scalp, and CSF probability maps, a subject-specific bimodal MR-CT neonatal head template is employed. In the next step, the subject-specific probability maps are fed to the expectation maximization algorithm in conjunction with Markov random field method implemented in FSL software to segment the skull and scalp from the input MR image. The results of the proposed method were evaluated through various experiments. First, we employed our method as a brain tissue extractor and compared its results with public methods such as the Brain Extraction Tool (BET) and Brain Surface Extractor (BSE). Second, we calculated the similarity in shape between the frontal and occipital sutures (which had been reconstructed from segmented cranial bones) and the ground truth. For this purpose, modified versions of the Dice similarity coefficient (DSC) were adopted and used. Finally, retrospective data including MRI and CT images obtained from the same neonate within a short time interval were used. After aligning the two images, the DSC and modified Hausdorff distance (MHD) were used to compare the similarity of the cranial bones in the MR and CT images. Furthermore, the anterior fontanel size was compared to the normal size reported for neonates in the same age range. Cranial bone thickness was calculated and compared to normal values reported for healthy neonates. The results of these experiments demonstrated the success of our segmentation method. The algorithm for creating subject-specific atlases is publicly accessible through a graphical user interface at medvispy.ee.kntu.ac.ir.

## 1. Introduction

Magnetic resonance imaging (MRI) has long been recognized as a valuable tool for studying neonatal brain development and injury due to its advantages of safety and relatively high spatial resolution [Nielsen et al., 2018; Gilmore et al., 2018; Li et al., 2019; Richter et al., 2022]. Brain extraction or skull stripping is a fundamental preprocessing step that segments an MR image into the brain and non-brain tissues, which is an indispensable and challenging task in neonatal neuroscience studies. This process is critical in brain tissue segmentation because under or over-estimation of brain tissue voxels cannot be salvaged in consecutive processing steps, potentially leading to error propagation through subsequent analyses (Serag et al., 2016; Gao et al., 2019; Wang et al., 2020).

Furthermore, skull segmentation can be used to provide a realistic head model for source localization using electroencephalography (EEG) or magnetoencephalography (MEG) signals (Okada & Delpy, 2003; Kiesler & Ricer, 2003; Li et al., 2015; Nielsen et al., 2018). Realistic MR imaging models of a neonate’s head are commonly used to identify different tissues with different electrical and optical properties in the head. Several studies have used realistic neonatal head models to investigate the effect of fontanels and sutures on EEG and MEG source analysis (Roche-Labarbe et al., 2008; Dehaes et al., 2013; Lew et al., 2013; Odabaee et al., 2014; Azizollahi et al., 2016; Antonakakis et al., 2019; Noreika et al., 2020). It is noteworthy that manual or semi-automatic skull segmentation was performed in most of the aforementioned studies because there is no fully automatic method available for skull segmentation from neonatal cerebral MR images. Manual or semi-automatic skull segmentation methods are very time-consuming and prone to intra- and inter-observer variability.

Despite the benefits of using MR imaging to study brain growth and identify, monitor, and control neuronal diseases, this modality is primarily helpful for soft tissue differentiation because cranial bones are poorly represented due to their low water content. This inconvenience is more pronounced in neonatal cerebral MRI compared to adults due to the fine and incomplete skull structure. The presence of higher motion artifacts and the newborn’s small head size are additional challenges in neonatal skull segmentation in MRIs. Additionally, higher-resolution scanning in neonates generates more noise and chemical shift artifacts (Weinreb et al., 1985; Prastawa et al., 2005; Cardoso et al 2013; Makropoulos et al 2018). Consequently, the neonatal skull appears in MR images with variable thickness and heterogeneous intensity, as opposed to the adult skull, which is represented as a thick and specific intensity structure. Accordingly, the skull segmentation methods developed for adult MRIs generally fail to perform accurately in neonates. On the other hand, neonatal head computed tomography (CT) is ideal for skull segmentation because it provides better contrast between the cranial bones and remaining tissues (Li et al., 2015). However, it fails to resolve cerebral soft tissues that are mandatory for neuroscience studies. Furthermore, due to the harmful ionizing radiation, CT scans can only be obtained from neonates when necessary for clinical purposes. (Mohtasebi et al.,2021).

Another major impediment to the development of accurate skull segmentation methods in neonatal cerebral MRIs is a lack of reliable ground truth(GT) for validating the results. Manual or semi-automatic skull segmentation performed on image slices by expert radiologists is mostly used to provide GT in cerebral MRIs of adults or infants (Gao et al., 2019). However, due to the aforementioned challenges, this technique is very time-consuming and impractical in cerebral MRIs acquired from neonates. To provide faithful landmarks and GT from three-dimensional MR data and smooth the path for developing accurate neonatal skull segmentation methods, new ideas inspired by anatomical knowledge and imaging technology are required. The intensity of an MR image in a region of interest is determined by the pulse sequence and tissue type. Some tissues with high water content, such as edema, tumors, infarction, inflammation, infection, and hemorrhage (hyperacute or chronic), appear dark on T1-weighted images (Keith A. Johnson). In contrast, some tissues such as fat, subacute hemorrhage, melanin, protein-rich fluid, slowly flowing blood, paramagnetic substances (gadolinium, manganese, copper), and laminar necrosis of cerebral infarction appear bright on T1-weighted images (Keith A. Johnson). In most cases, calcification of tissues has no effect on MR images or appears as a reduction in signal intensity and darkening of the tissue (Keith A. Johnson). Henkelman et al. (Henkelman et al., 1991) demonstrated that it is displayed brightly and at high intensities in some cases. In neonatal T1-weighted MRIs, some sutures are visible as bright spots in specific slices due to ossification. The bright spots can be distinguished via intensity filtering and examining the adjacent slices along with the three-dimensional views and used as anatomical landmarks for validation of the skull segmentation results. Another method of validation is to compare the results of skull extraction from MR and CT images acquired within a short time interval from the same subjects. Obviously, such bimodal data from neonates with anatomically normal heads are rarely available in medical archives because they can only be obtained for clinical purposes. In this regard, Nielsen et al. (Nielsen et al., 2018) assessed the accuracy of skull extraction results from MR images by comparing CT-based skull segmentations on a group of adult subjects.

Despite the importance of accurate scalp and skull segmentation in neonatal MR images (Wang et al., 2020), there are few published methods in this field, owing to the aforementioned difficulties. In this context, Burguet et al. (Burguet et al., 2004) created realistic models of children’s heads from 3D-MRI segmentation using a watershed algorithm. Despotovic et al. (Despotovic et al., 2009) extracted scalp, fat, skull, cerebrospinal fluid (CSF), and brain tissues from a 39-week-GA newborn T1 image using a hybrid algorithm that included active contours, fuzzy C-means clustering, and mathematical morphology. Daliri et al. (Daliri et al., 2010) extracted skulls from infants’ MRI using a Hopfield neural network and a fuzzy classifier. Yamaguchi et al. (Yamaguchi et al., 2010) suggested a fuzzy-logic-based method for skull stripping in children’s MR images. They proposed an active surface model-based skull-stripping method. Within a Bayesian classification scheme, the intensity is modeled using a Gaussian mixture model. Prior information is derived from tissue probability maps in a built atlas. To extract the brain region, P’eport’e et al. (Péeportée et al.,2011) developed an atlas-free segmentation method based on morphological operations, region growing, and edge detection. Adaptive thresholding is performed on each 2D slice separately using a threshold derived from k means clustering. Mahapatra et al. (Mahapatra, 2012) used prior shape information within a graph cut framework and a smoothness term based on gradient information for skull stripping of neonatal cerebral MRI. Similarly, Kobashi and Udupa (Kobashi & Udupa, 2013) presented an active surface method using a prior fuzzy shape model for skull segmentation. To improve skull removal from pediatric MR images, Shi et al. (Shi et al., 2012) developed a brain-extraction meta-algorithm and integrated it into the analysis and neonatal brain extraction toolbox (iBEAT). A level-set based segmentation algorithm is used to fuse and refine brain extraction using different parameters. Shi et al. (Shi et al., 2012) use the affinity propagation technique to select atlases based on their intensity similarity to reduce computation complexity. Serag et al. (Serag et al.2016) developed a brain segmentation method called accurate learning with few atlases (ALFA), within a multi-atlas segmentation framework. It performs classification of the MRI into the brain and non-brain regions using either a naive Bayes or linear discriminant analysis classifier. Atlas selection is performed to identify atlases that provide complementary information across an atlas database. They used subject-specific atlases of manually segmented images in their database and used individual images as reference. Noorizadeh et al. (Noorizadeh et al., 2019) presented a multi-atlas patch-based label fusion method for automatic brain extraction from T2-weighted neonatal MR images.

Most of the aforementioned methods suffer from unrealistic and low accuracy rates of neonatal skull segmentation results. This low performance is expected and comprehensible for the methods that use low-level image processing tools (Burguet et al., 2004 ; Despotovic et al., 2009; Yamaguchi et al., 2010 ; Péeportée et al 2011) because they rely on image intensities to discern between tissues, whereas the skull is quasi-imperceptible in neonatal MR images. Artificial intelligence-based and atlas-based methods (Daliri et al., 2010; Mahapatra, 2012; Noorizadeh et al., 2019; Shi et al., 2012), on the other hand, yielded unsatisfactory results mainly due to the imperfection of their skull geometry knowledge -base. Serag et al. (Serag et al.2016) compared their method with eleven other brain segmentation techniques, including the Brain Extraction Tool (BET) (Smith, 2004) and the Brain Surface Extractor (BSE) (Shattuck et al., 2001).

The tissue of interest in skull stripping methods is the brain, and to the best of our knowledge, skull segmentation was mostly ignored. In the current study, we have developed develop a new method for extracting the skull, scalp, and intracranial tissues from neonatal T1-weighted MRI. The main methodological contribution of this paper over its predecessors is taking advantage of a bimodal MR-CT atlas and a realistic and accurate skull model created using a retrospective CT dataset. Inspired by (Cardoso et al., 2013; Cherel et al., 2015; Serag et al., 2016), subject-specific probability tissue maps are generated based on the MRI and CT atlas population in the age range of 39 to 42 weeks gestational age (GA). The probability maps of the skull and scalp are constructed from CT images and used as a priori information alongside the CSF probability map to extract the skull and fontanels from MR images. The adaptation of this model and the input MR image to be segmented is the advantage of this method. In other words, having this bias of the template relative to the input image helps to improve segmentation results. For skull extraction, the EM algorithm is then used in conjunction with the Markov random field (MRF) implemented in FSL software (FAST toolbox) (Smith et al., 2004) to which the created subject-specific probability tissue maps are fed. The extracted skull is then used to obtain intracranial tissue. An overview of the proposed methodology is shown in Figure 1. Our proposed method to generate subject-specific scalp, skull and CSF probability maps has been integrated into MedVisPy, a medical desktop software application written in Python, to support clinicians in processing, analyzing, and visualizing medical images. It can be downloaded from medvispy.ee.kntu.ac.ir. The probability maps generated by MedVisPy are subsequently fed to FSL for extracting scalp, skull and intracranial from a neonatal cerebral MR image.

**Fig. 1.**
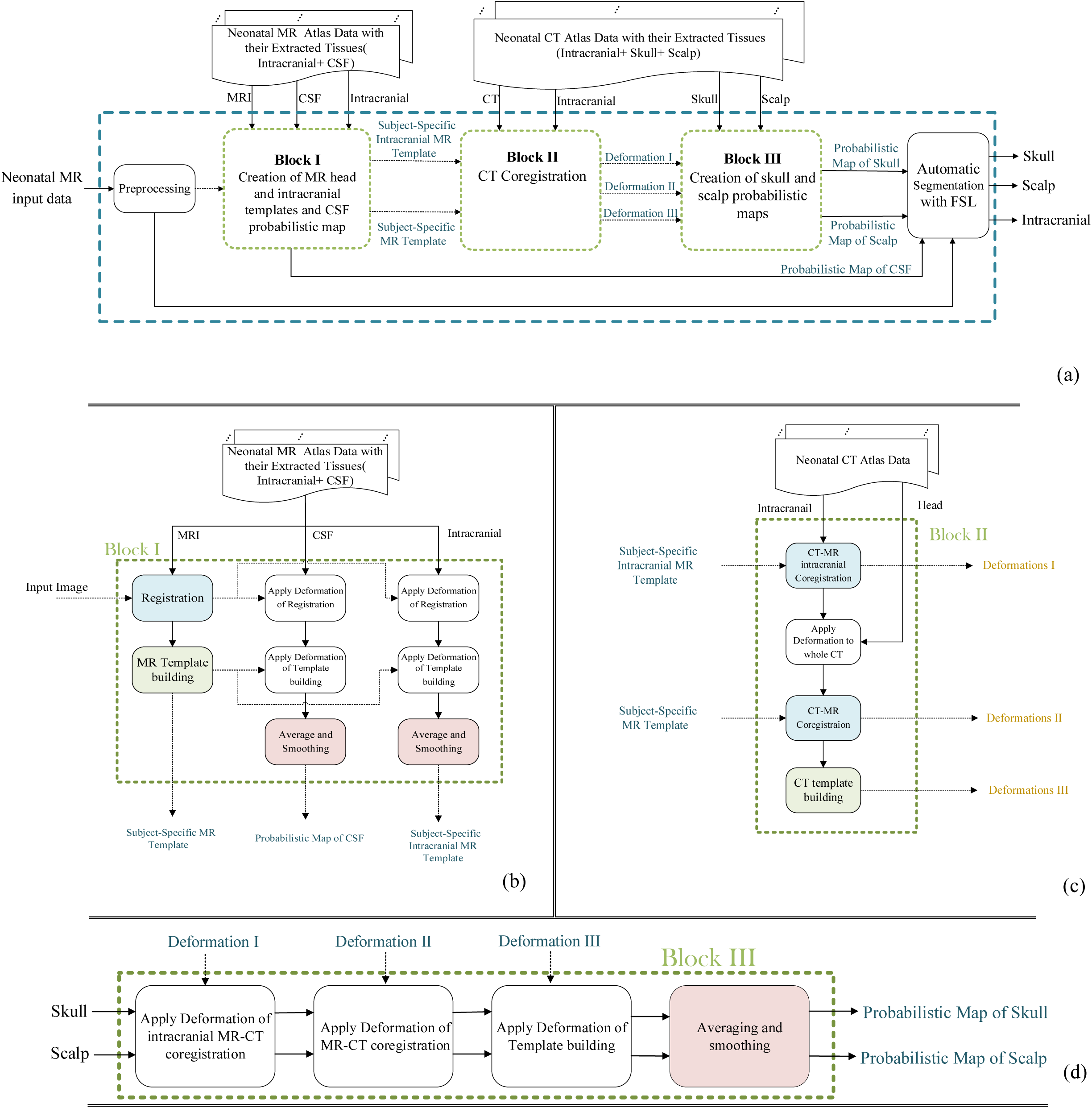
a) The proposed pipeline to extract scalp, skull and intracranial from an input neonatal cerebral MR image. b) The details of Block I. c) The details of Block II. d) The details of Block III

## 2. Materials and methods

### 2.1 Subjects and Data Acquisition

In our retrospective study, two databases of MR and CT images were used. All images were examined by our expert radiologist and no serious cerebral anatomical abnormality was detected. The MRI database was employed to construct geometric and probability models of newborn heads. The CT dataset was also used to build a geometric model of the neonatal head and probability models of the skull and scalp. In the following, these two datasets are introduced, respectively.

#### MRI database

The MRI data employed in this study were exactly the same as those used in (Kazemi et al., 2007). The subjects in the MRI database were 14 newborns of GA between 39 and 42 weeks at the date of MR examination and were partitioned into two subsets. The first subset consisted of 7 subjects, including 4 girls and 3 boys, which were used to create a geometric model. These images have also been used as test images within a leave-one-out cross-validation strategy. The second subset consisted of 7 MRIs, including 3 girls and 4 boys, that were used only for evaluating skull segmentation results. Twelve newborns were imaged by a General Electric 1.5T machine using the following sequence parameters: TR=10.1 ms, TE=2.208 ms, TI=500 ms. Each volumetric image had 512 × 512 pixels per slice with a 220 mm field of view with a voxel size of 0.47× 0.47× 0.47 mm^3^.

The two remaining images were obtained by a Siemens 3 T MR scanner with the following sequence: TR=1820 ms, TE=4.38 ms, and TI=1100 ms. Each volumetric image had 256 × 256 pixels per slice with a voxel size of 1×1×1 mm^3^. Then, to reduce the computation time, all images were resliced to 0.94 × 0.94 × 0.94 mm^3^ using the trilinear interpolation method in SPM software.

An ideal dataset for the evaluation of our skull segmentation method is a bimodal MR-CT image acquired simultaneously from a normal subject. Such ideal data being rarely available, we used a CT image of a subject with the age of 40 weeks GA and an MR image of the same subject at 42 weeks GA. The CT and MR image resolutions were 0.32 × 0.32 × 0.63 mm^3^ and 0.5 × 0.5 × 0.5 mm^3^, respectively.

#### CT database

The images used in this study were exactly the same as those used to build the CT atlas in (Ghadimi et al., 2015; Ghadimi et al., 2017; Mohtasebi et al., 2021). The dataset contains 16 CT images, including 12 boys and 4 girls of infants aged from 39 to 42 weeks at the date of examination (GA).

All images were acquired using a LightSpeed 16, GE Medical Systems. The imaging matrix was 512 × 512 pixels. The voxel size was between [0.26-0.49] × [0.26-0.49] × [0.6-1.25] mm^3^ with a median of 0.32 × 0.32 × 0.63 mm^3^. Similar to MRIs, all non-axial images were reoriented to the axial plane and resliced to 0.94×0.94×0.94 mm^3^ isotropic voxels. There was no subject in common between MR and CT databases that have been used for geometric and probability model creation.

### 2.2 Atlas Data Preparation

The neonatal skull segmentation algorithm, as shown in Figure 1, receives three inputs: neonatal MR input data, MR atlas data including raw images and their extracted intracranial and CSF tissues, and CT atlas data including raw images and their extracted scalp, skull, and intracranial tissues.

#### 2.2.1 MR Atlas Data Preparation

Neonatal MR atlas data provide a part of the anatomical a priori information for the skull segmentation algorithm. The MR atlas images were semi-automatically preprocessed before being fed into the algorithm. The neck and extra parts of the head were removed manually using 3D Slicer software(Fedorov et al., 2012). Then, noise elimination, rotation, bias correction, and histogram matching were performed. The bias correction was done by the N4 algorithm (Tustison et al., 2010). Finally, the images were smoothed using a 2 mm Full Width at Half Maximum (FWHM) Gaussian kernel. The intracranial and CSF tissues have already been segmented automatically followed by manual corrections (Kazemi et al., 2008) and have been examined by our expert radiologist, and are available in the database.

#### 2.2.2 CT Atlas Data Preparation

CT atlas data complement the anatomical a priori information for the neonatal skull segmentation method. Before being fed into the skull segmentation algorithm, CT atlas images were subjected to a number of preprocessing steps. MRIcron software was used to remove the neck and extra parts of the head first. In addition, some unwanted structures visible in some CT images, such as pacifiers, masks on the infant’s mouth, tubes, neck, and noise around the image, must be manually removed. To solve the aforementioned issue, the head mask of the input raw image must be extracted. To accomplish this, we used the histogram thresholding (Otsu, 1979) method and morphology operators such as opening, closing, filling, and identifying connected components to separate the head from the background on the input raw image. Then, by multiplying the head mask by the CT image, everything inside the head kept its intensity while everything in the CT image’s background received zero intensity. CSF and the background have the same intensity in CT images, which is zero. To differentiate between CSF and background in CT images, we assigned a -1000 intensity to each voxel in the background. The intensity of the CT images was then transformed from the Hounsfield unit to the Cormack unit using reversible mapping proposed by Rorden et al. (Rorden et al., 2012). This intensity transformation increases the dynamic range of soft tissues, gets the intensity of CT images closer to the intensity of MR images, and improves registration accuracy.

After pre-processing, the intracranial, scalp, and skull tissues must be extracted from the CT atlas population. The method proposed by Ghadimi et al. (Ghadimi et al., 2017) was used to extract intracranial tissue from CT atlas images. To begin, the coupled level sets algorithm (Ghadimi et al., 2015) was used to determine the inner surface of the skull. The resulting surface (cranial bone inner surface and sutures) was assumed to be the outer surface of intracranial tissue. After manual correction by a specialist, a mask was created by the filling operation and employed to extract the intracranial from the CT image. Figure 2 depicts the process of extracting intracranial tissue from a CT image. The skull was extracted using Otsu’s method of histogram thresholding (Otsu, 1979).

**Fig. 2.**
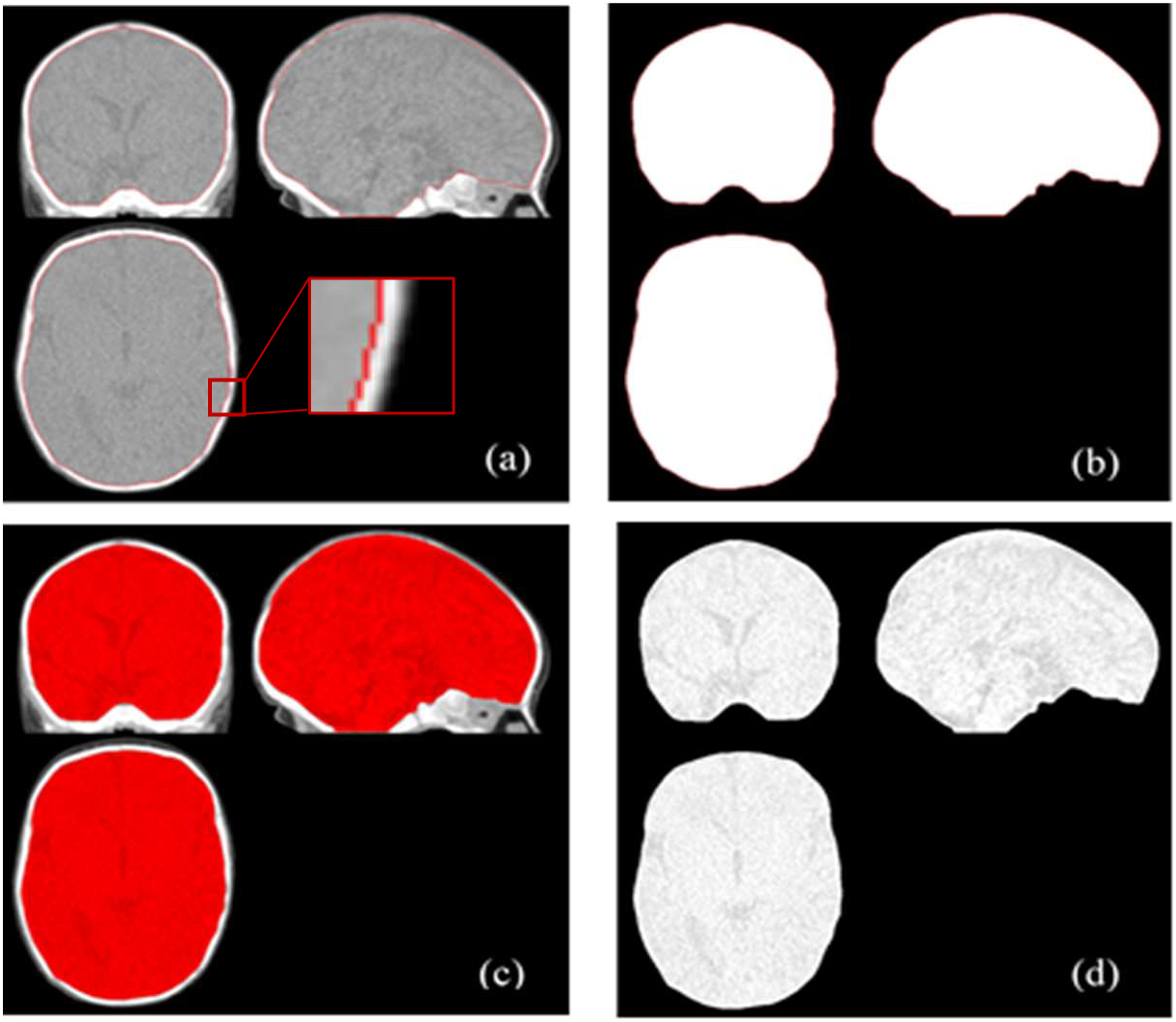
Extraction of CT intracranial tissue using the method proposed in (Ghadimi et al., 2015) a) The extracted inner surfaces of cranial bones b) Corresponding mask of the resulting inner contour c) The extracted mask overlaid on the CT image d) The final extracted intracranial

The fontanels and sutures, which are the intersection of inner and outer skull contours, were reconstructed using the coupled level sets algorithm (Ghadimi et al., 2015) to extract the scalp from the CT images. After manual correction in some cases, we used a logical OR operation between the skull and the reconstructed fontanels to create a continuous volume of the skull. The coupled level sets algorithm (Ghadimi et al., 2015) was then used to determine the outer surface of this volume. The filling operation produced a mask of this closed surface. Finally, this mask was used to remove all areas inside the skull, including the cranial tissue. Figure 3 shows the extraction of scalp tissue from a CT image.

**Fig. 3.**
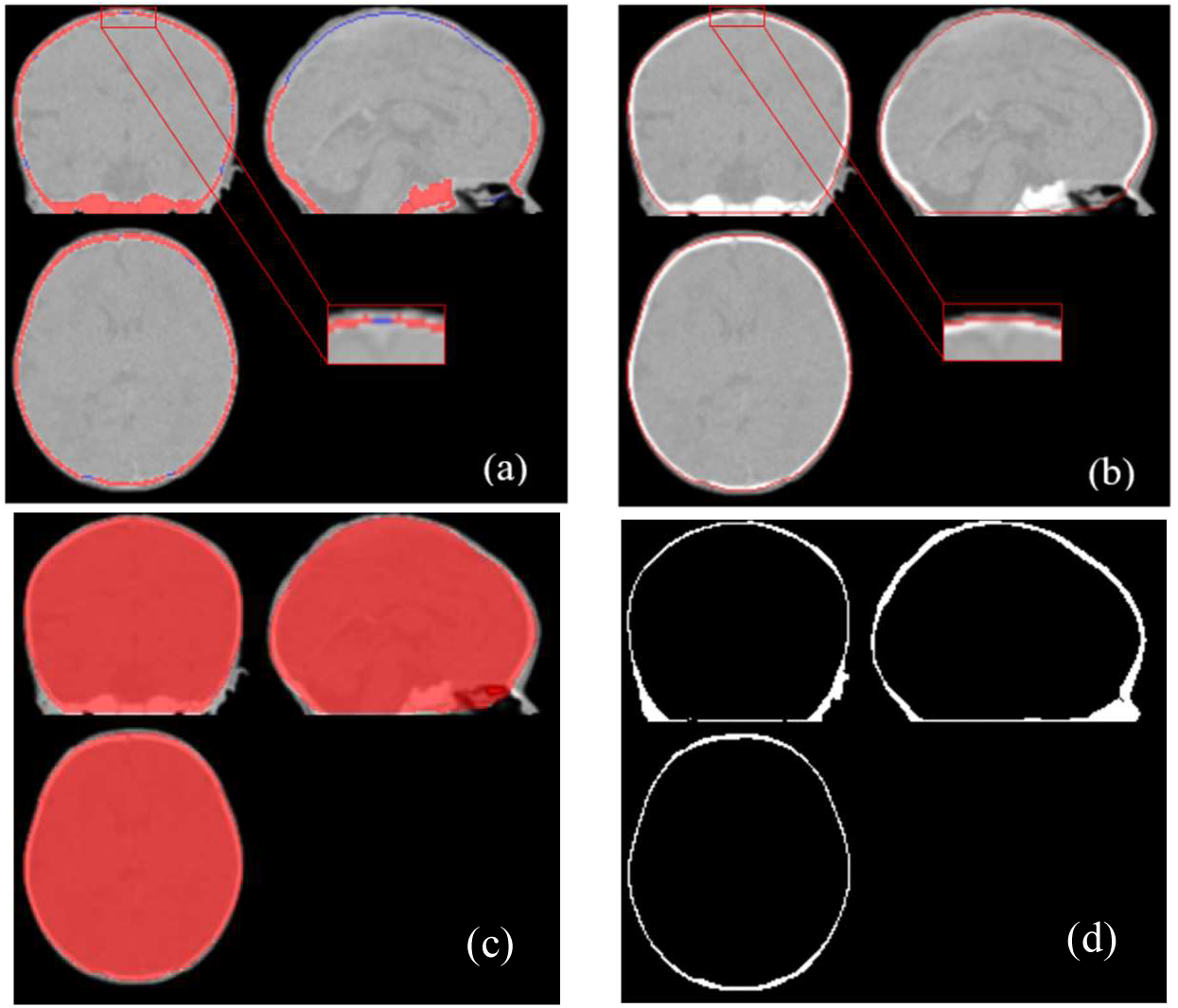
Extraction of CT scalp tissue. a) The reconstructed fontanels (blue) with cranial bones (red) overlaid on the CT image b) Outer contour of the obtained closed volume of the skull c) The corresponding mask overlaid on the CT image d) The final extracted scalp

### 2.3 Skull Segmentation Algorithm

#### 2.3.1 Preprocessing of the Input MR Image

The skull segmentation algorithm receives a neonatal cerebral MR image as input. As shown in Figure 1, this image was preprocessed to be prepared for further processing. The preprocessing stage was explained in Section 2.2.1.

#### 2.3.2 Creation of Subject-Specific MR Atlases

The MR atlas images in the database were mapped to a reference space, as shown in Figure 1(b) (Block I in Figure 1), to construct a subject-specific geometric model of the head as suggested by (Cardoso et al., 2013; Cherel et al., 2015; Serag et al., 2016). Accordingly, the MRI of the subject to be segmented was chosen as a reference, and each subject in the MR atlas population was registered to it using a rigid, followed by a non-rigid transformation. In the first step, a 12-parameter affine transformation (including translation, rotation, scaling, and shearing) was used to align the MR atlas population with the reference space. The linearly mapped MR images were then matched to the reference using a nonlinear symmetric normalization (SyN) transformation (Avants et al., 2008) and using mutual information (MI) similarity measure. All deformations were applied with the help of ANTs software, which was developed within the framework of the ITK open-source tool (Avants et al., 2014). Furthermore, the registration parameters were applied to the extracted tissues of each MR image in the database, including intracranial and CSF. The goal was also to align these tissues with the desired reference.

Subsequently, an MRI template was created using a groupwise registration approach (Avants et al., 2014). This method is available in ANTs software via the script “antsMultivariateTemplateConstruction.sh”. It was run for two iterations, with cross-correlation (Lemieux et al., 1998) used as a similarity index and the SyN energy term used to build a subject-specific MRI template.

The mapped intracranial and CSF tissues were then transformed into the MRI template’s space using the deformation parameters obtained during template construction. It is worth noting that the CSF tissue was extracted in binary format. Finally, the tissues were averaged and smoothed using a Gaussian filter with an FWHM of 2 mm. An intracranial geometric model and a CSF probability model were created in this manner. A tissue probability atlas is a 3D image with the same size as the MRI (or CT) template. The value of each voxel 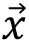 in the probability atlas represents the probability of the presence of the *i*th tissue 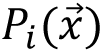 in the same voxel of the corresponding template. Figure 4(c) illustrates the intracranial MRI template developed in this section for segmenting the subject. Similarly, Figure 4(d) portrays the CSF probability map created for the same subject.

**Fig. 4.**
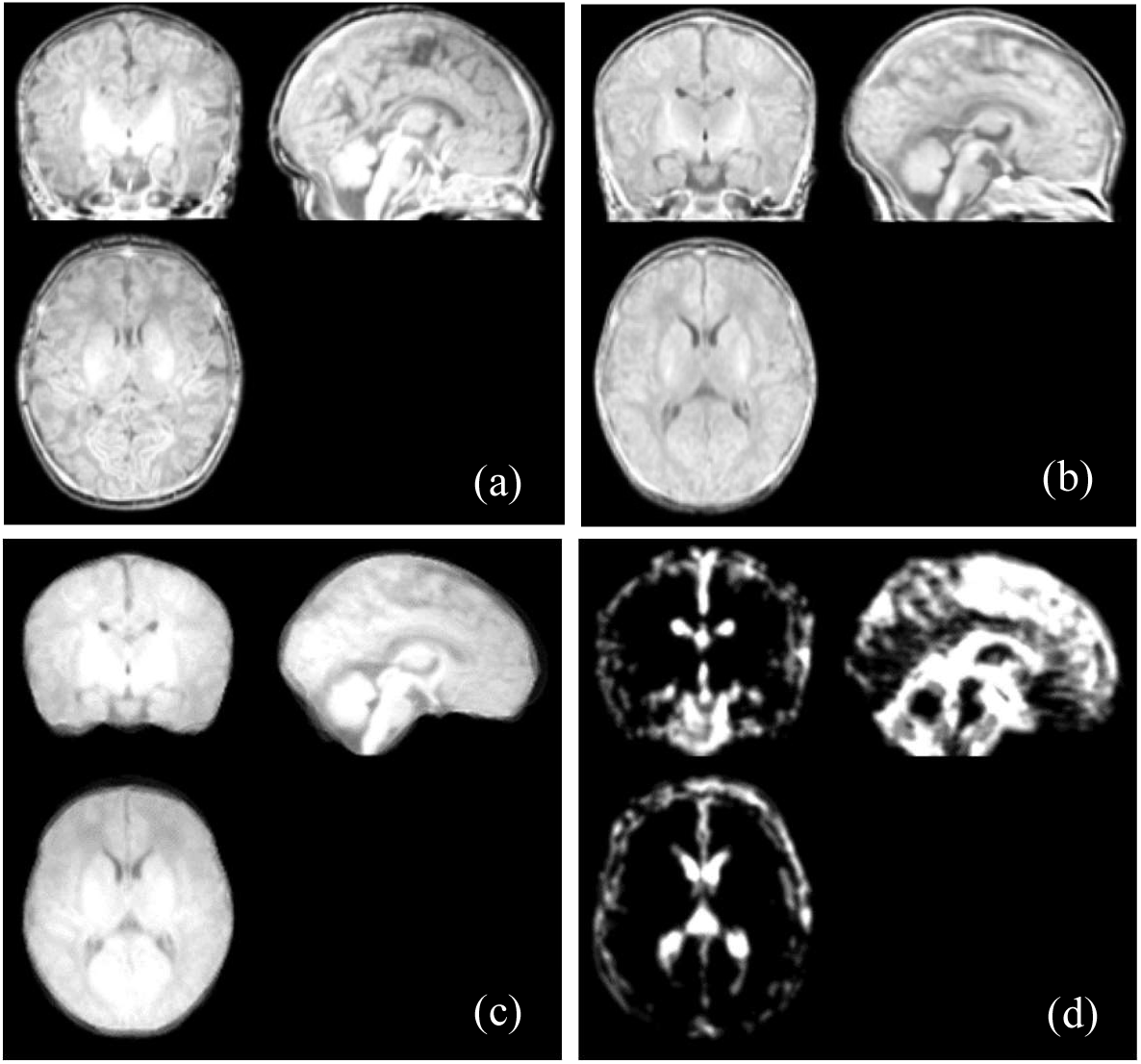
Subject-specific MR templates for subject N3 a) MRI of subject N3 b) Whole head MR template c) Intracranial MR template d) CSF probability model

#### 2.3.3 Creation of Subject-Specific CT Atlases

As shown in Figure 1(c) (Block II in Figure 1), the method proposed by Ghadimi et al. (Ghadimi et al., 2017) was used to construct a subject-specific CT template corresponding to the input MRI. The only difference between our method and theirs was the reference space used to map the images. In this study, the geometrical templates made in Section 2.3.2 served as the reference space. As a result, in order to segment each input MRI, all atlas CT images had to be mapped to the new subject-specific space represented by the full head and intracranial MRI templates created in Section 2.3.2.

##### CT-MR Co-registration

The mapping of atlas CT images to the subject-specific full head and intracranial MRI templates consisted of three main steps:

1. First, using 12-parameter affine and SyN transformations, the atlas CT intracranial images were linearly and nonlinearly transformed to the subject-specific intracranial MRI template, respectively (Avants et al., 2008). The CT intracranial images were matched to a subject-specific MRI intracranial template using MI.
2. Second, the linear and nonlinear transformation matrices derived from mapping each atlas CT intracranial image were applied to its original whole head image.
3. Third, using an affine transformation and the MI index, the resulting whole-head CT images were registered to the subject-specific MRI template.

##### Subject-specific CT template building

The CT atlas images that had been successfully mapped to the subject-specific MRI template were employed to construct the subject-specific CT template at this stage. The procedure was similar to that described in Section 2.3.1 for creating an MRI template. Figure 5 shows a CT template built for one of the MRI test subjects.

**Fig. 5.**
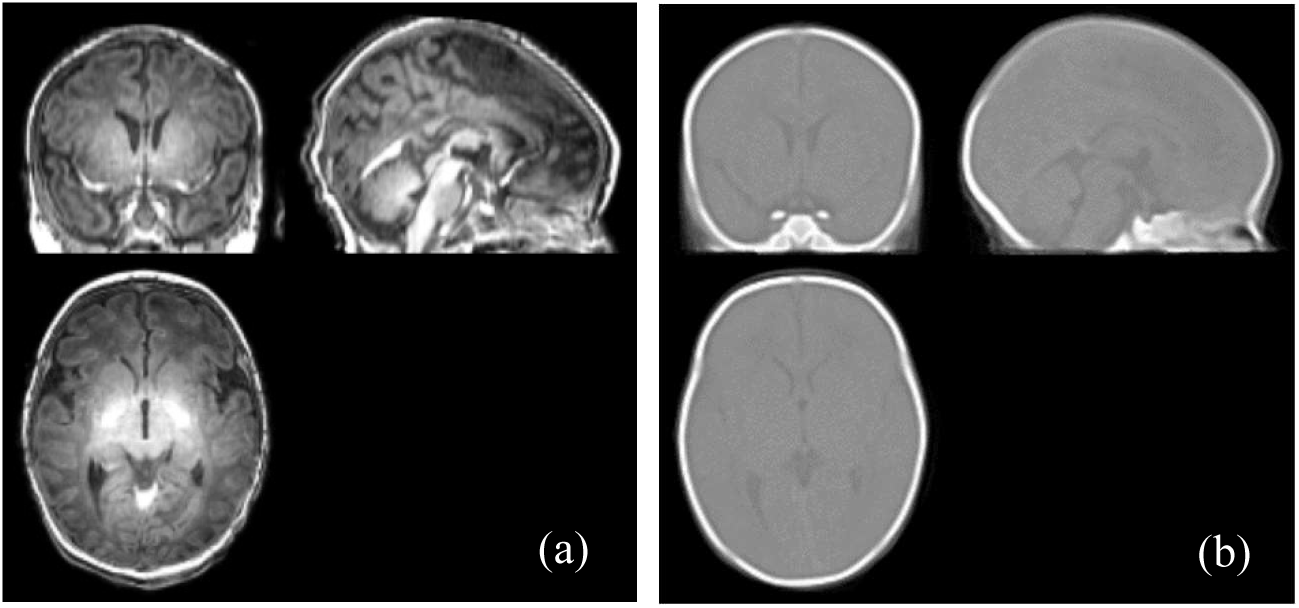
a) MRI of subject T6. b) The specific CT template for this subject

#### 2.3.4 Creation of Subject-Specific Probability Maps of Neonatal Skull and Scalp

The atlas CT skull and scalp images prepared in Section 2.2.2 (CT atlas data preparation) were used to create probability maps of the skull and scalp tissues (see Figure 1(d), Block III in Figure 1). First, the CT atlas skull and scalp images were transformed using the transformation parameters obtained during CT-MR co-registration. The goal was to align these tissue images with the subject-specific MR template. Second, using the deformation parameters obtained during CT template construction, the mapped skull and scalp tissues were transformed to the CT template’s space. Finally, the mapped images of the tissues were averaged and smoothed with a Gaussian filter at FWHM=2mm. Figure 6 depicts the subject-specific probability maps of the skull and scalp constructed for one of the MR test subjects.

**Fig. 6.**
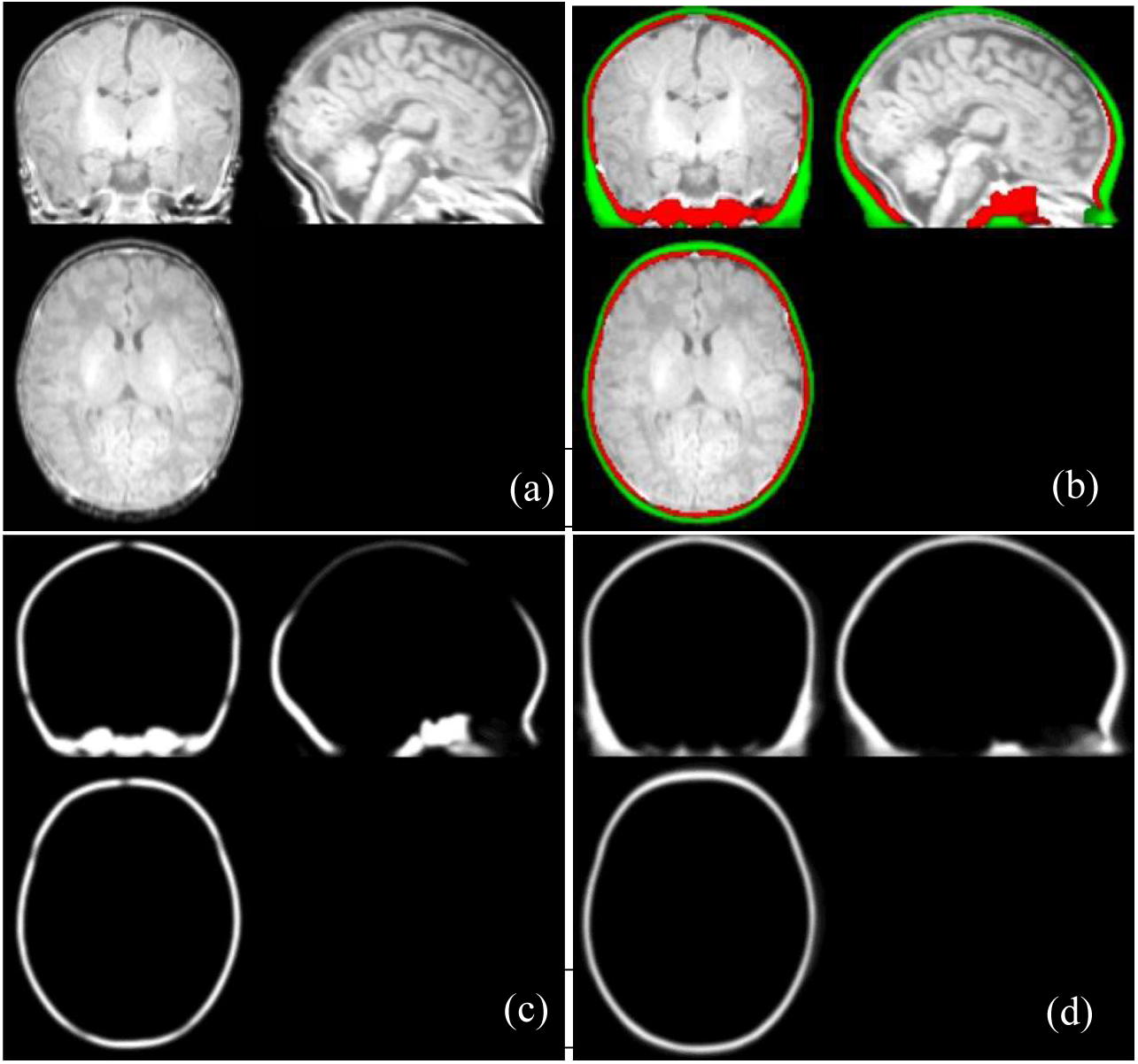
a) MRI of subject N2 b) The subject-specific probability models overlaid on subject N2. c) The specific probability model of neonatal skull for this subject d) The specific probability model of neonatal scalp for this subject

#### 2.3.5 Skull Segmentation from the Input MR Image

After creating subject-specific probability maps for the skull, scalp, and CSF (Sections 2.3.4 and 2.3.2), the skull can be segmented from the preprocessed input MR image using the FAST toolbox of conventional FSL software (Smith et al., 2004). Segmentation was also performed using subject-specific neonatal probability maps created from MR and CT atlas images (instead of using the FSL default models). The identity matrix was used to map the input image to the probability models because the preprocessed input image and the subject-specific probability maps were in the same space. In the probability maps, probabilities less than 0.4 were considered as zero. This step produced binary masks of the skull, scalp, and intracranial, as shown in Figure 7(a). Histogram thresholding was used to extract the binary skull mask (Figure 7(b)). In some cases, post-processing of the extracted skull was required, such as noise removal and removal of some pixels misidentified as skull tissue.

**Fig. 7.**
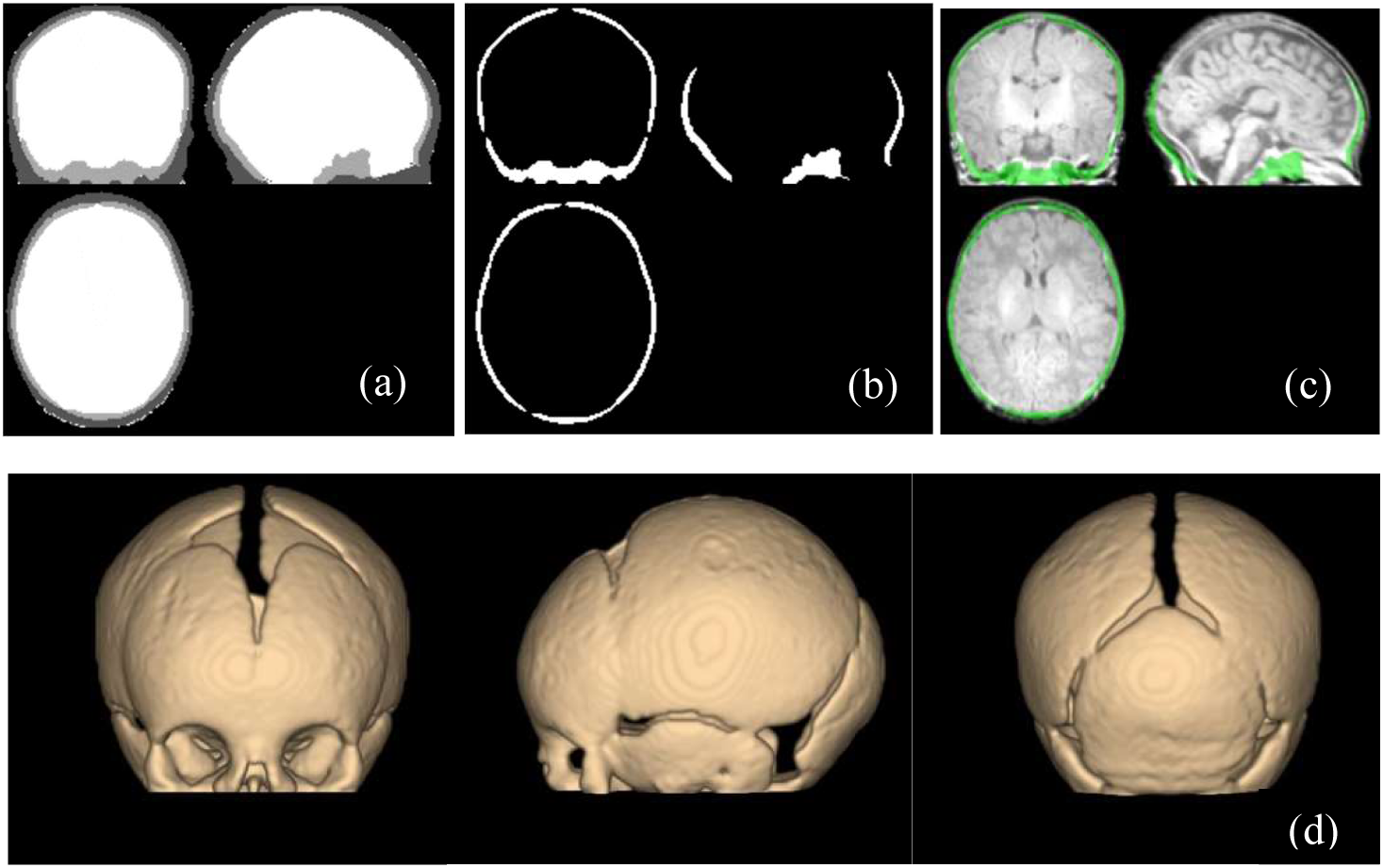
a) Results of the FSL software. b) The final extracted skull for subject T2 c) Extracted skull overlaid on MRI d) Three-dimensional rendering of the segmented skull

### 2.4 Evaluation Methods

GT is required to validate the results of skull segmentation from neonatal MR images. It is frequently created by an expert using manual image segmentation. However, due to the lack of reliable intensity information for the skull, sutures, and fontanels in 2D slices, manual segmentation of cranial bones in neonatal MR images is a too complex and time-consuming task. As a result, anatomical landmarks on the 3D surface of the head in the MR image are used to evaluate the proposed method (Li et al., 2015). An alternative commonly practiced approach is to extract intracranial tissues after skull segmentation and compare them to manual intracranial segmentation as GT. Finally, since cranial tissues can be easily extracted in CT scans, CT images can be used as the required GT in subjects whose MR and CT images are available within less than two weeks of each other. In this section, three scenarios are presented to evaluate the results of the neonatal skull segmentation method.

#### 2.4.1 Evaluation of the Results Using the Intracranial Tissue Extraction

The first scenario for evaluating the results is to extract intracranial tissues from MRIs and compare them to GT. As stated in Section 2.2.1, the intracranial tissues are available for seven atlas MR images. They can be used as GT to evaluate our results within a leave-one-out framework.

To extract intracranial tissues, as explained in Section 2.3.5, histogram thresholding was used. In some cases, post-processing of the extracted intracranial was required, such as noise removal and removal of some pixels misidentified as intracranial tissue. The Dice similarity coefficient (DSC), sensitivity, and specificity were calculated using (1)-(3), respectively. They were also compared to the results obtained by commonly used and freely available programs such as BET (Smith, 2002), and BSE (Shattuck et al., 2001).

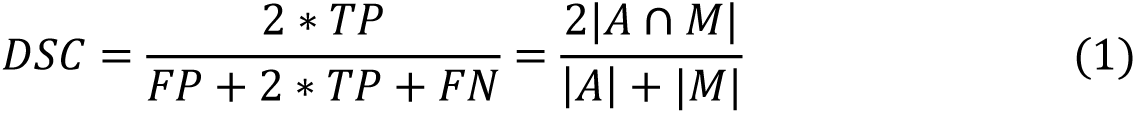

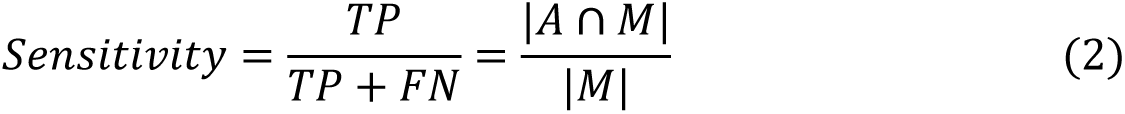

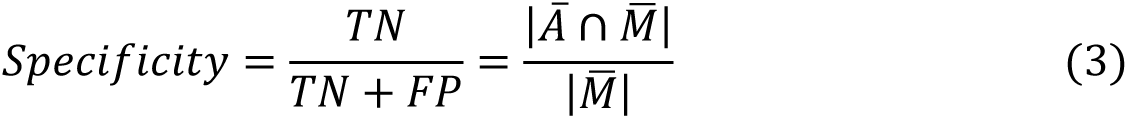

DSC represents the relative overlap between segmented tissue (A) and corresponding GT (M). Also, TP, FP, TN and FN represent the true positive rate, false positive rate, true negative rate, and false negative rate, respectively Higher values of DSC (Dice, 1945) indicate better skull stripping results. The algorithm’s sensitivity describes how sensitive it is to brain tissues. It indicates the likelihood of brain tissues being correctly identified. Another aspect is specificity, which describes the likelihood of correctly identifying non-brain tissue. To generate mean maps of these metrics, we averaged the FP, FN, and absolute errors across subjects. In addition, the modified Hausdorff distance (MHD) is commonly used to assess the spatial consistency of the overlap between two binary images by calculating the mean of the distances (ℎ_mean_) between them (Dubuisson & Jain, 1994 ; Ghadimi et al., 2015). It is given by:

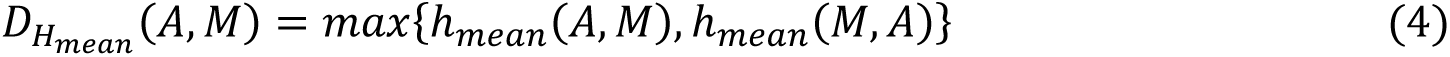

where

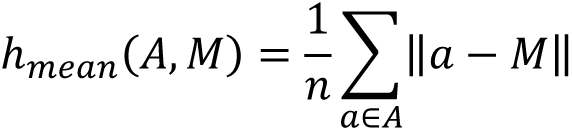

#### 2.4.2 Evaluation of the Results Using 3D Anatomical Landmarks

Figure 8 depicts the appearance of metopic and coronal sutures in the axial view of a T1-weighted MR image. The zoomed-in parts in Figure 8 illustrate these anatomical landmarks better. In this study, MRIcroGL software was employed by an expert to label these bright spots as sutures in 2D axial slices. For this purpose, the desired anatomical landmarks in each slice were first designated by drawing a square around them. Then, the pixels with high intensity in each square were labeled as suture. As different tissue types have different image intensities, thresholding the intensity and changing transparency in 3D volume rendering can reveal the desired tissues while making other tissue types transparent. Figure 9 (a) depicts the sutures as bright tracks in the 3D view of the head. The voxels in Figure 9 (b) were labeled as sutures using the aforementioned method. It is worth noting that all extracted anatomical landmarks were approved by an expert radiologist (MG).

**Fig. 8.**
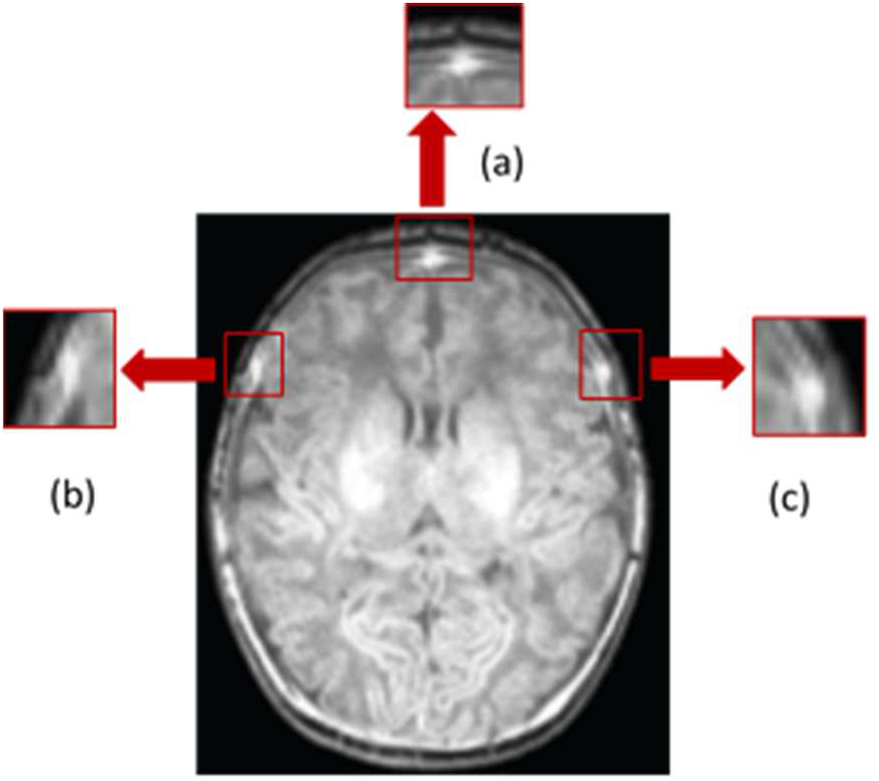
Axial view of the MR image of the subject N1. The bright spots enclosed by red rectangles show how sutures appear in a T1-weighted MRI. a) Metopic suture b, c) coronal sutures.

**Fig. 9.**
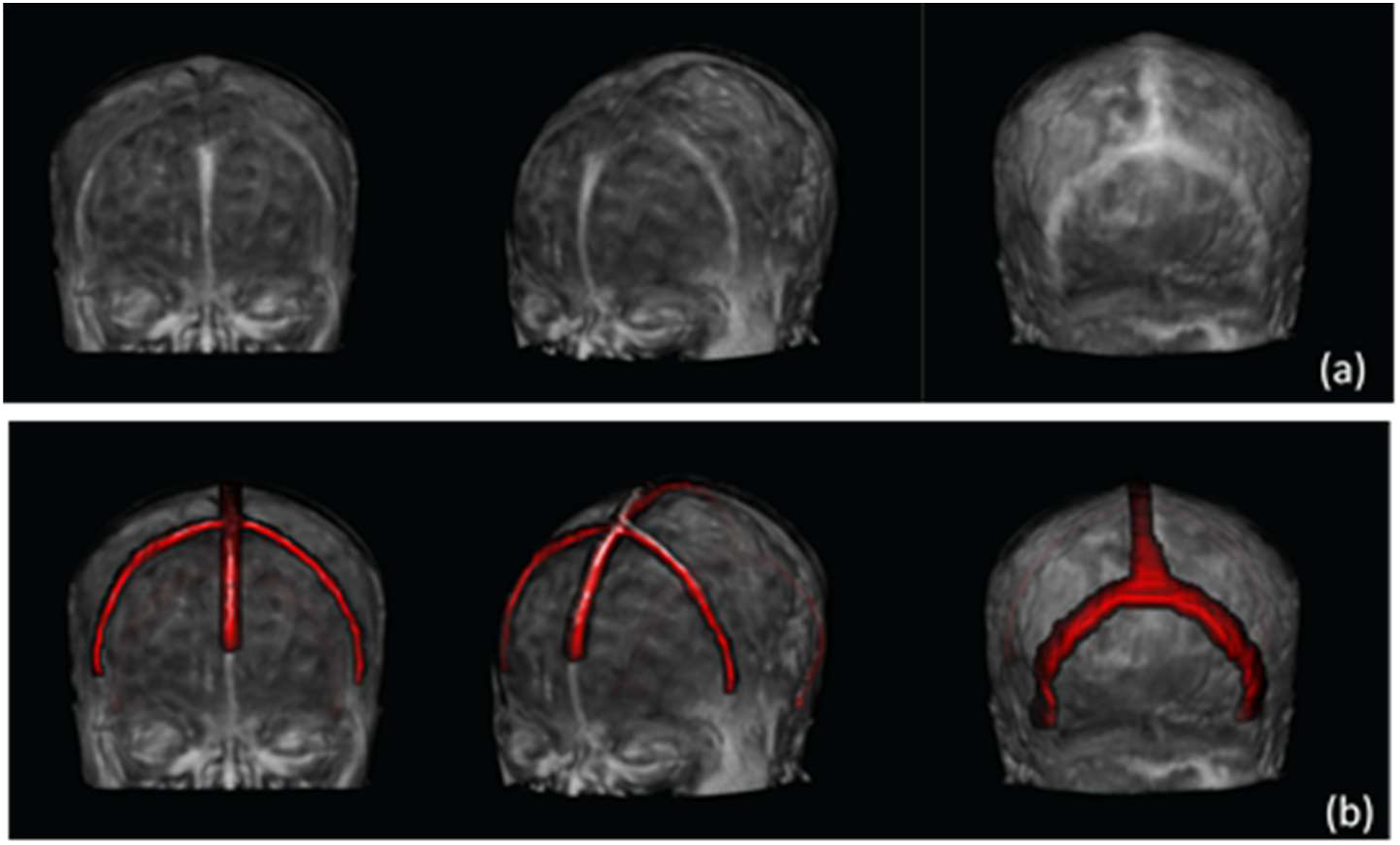
Manual extraction of sutures from MR image of subject N3. a) The sutures are visible as bright tracks in the 3D view of the head, b) Overlay of extracted voxels as anatomical landmarks in red with MR image.

The accuracy of the extracted sutures in the MR images was assessed qualitatively and quantitatively by examining the correspondence between their position and the position of extracted cranial bones. The position of manually extracted sutures on MR axial slices of a test subject is shown in Figure 10.

**Fig. 10.**
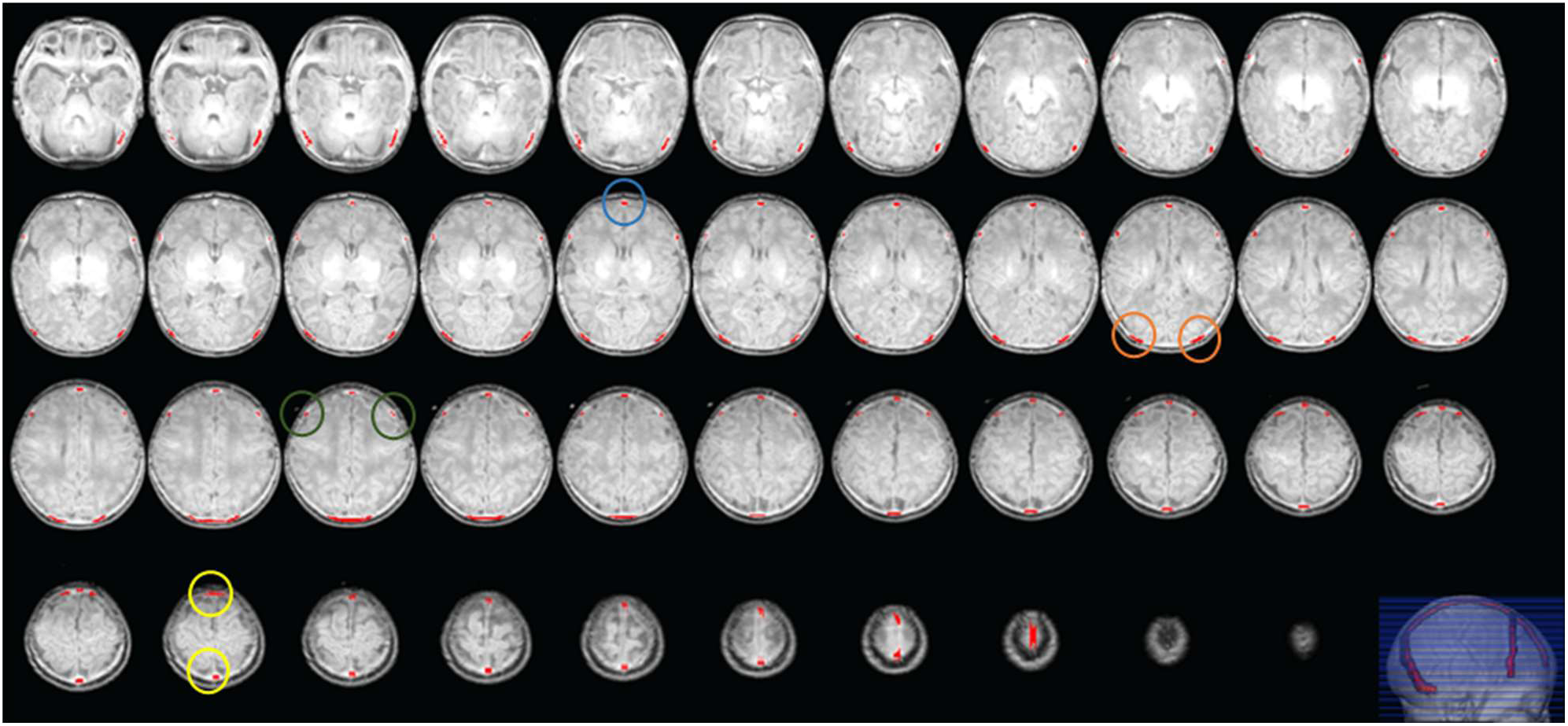
Axial MR slices of the subject N3. Red tracks in each slice show the anatomical landmarks corresponding to manually extracted sutures. The blue, green, orange and yellow circles enclose a metopic, coronal, lambdoid and sagittal suture, respectively.

For quantitative evaluation, we first used the coupled level set algorithm to reconstruct the fontanels from the extracted cranial bones (Ghadimi et al., 2015). The shape similarity between the GT landmarks and reconstructed fontanels was then calculated. The DSC is almost used for this purpose. However, both the manually extracted landmarks and automatically reconstructed fontanels are narrow and thin structures distributed three-dimensionally as a complementary part of the neonatal cranial bone. This is why DSC does not exactly match our requirements for this specific evaluation problem. Therefore, we defined and used a modified similarity index (MSI) as the number of corresponding landmark and fontanel voxels divided by the number of landmark voxels. Strictly speaking, in both landmark and fontanel images, two correspondent voxels must have the same coordinates.

However, by broadening the correspondence to the first and second nearest neighbors, two MSI variants, MSI-1 and MSI-2, were introduced and used in this study. MSI, MSI-1 and MSI-2 were calculated using Algorithm 1. Figure 14 depicts the qualitative match between anatomical landmarks (anterior and posterior sutures) and fontanels reconstructed by the coupled level set algorithm (Ghadimi et al., 2015). Both structures were superimposed on cranial bones segmented by the proposed algorithm.

**Algorithm. 1.**
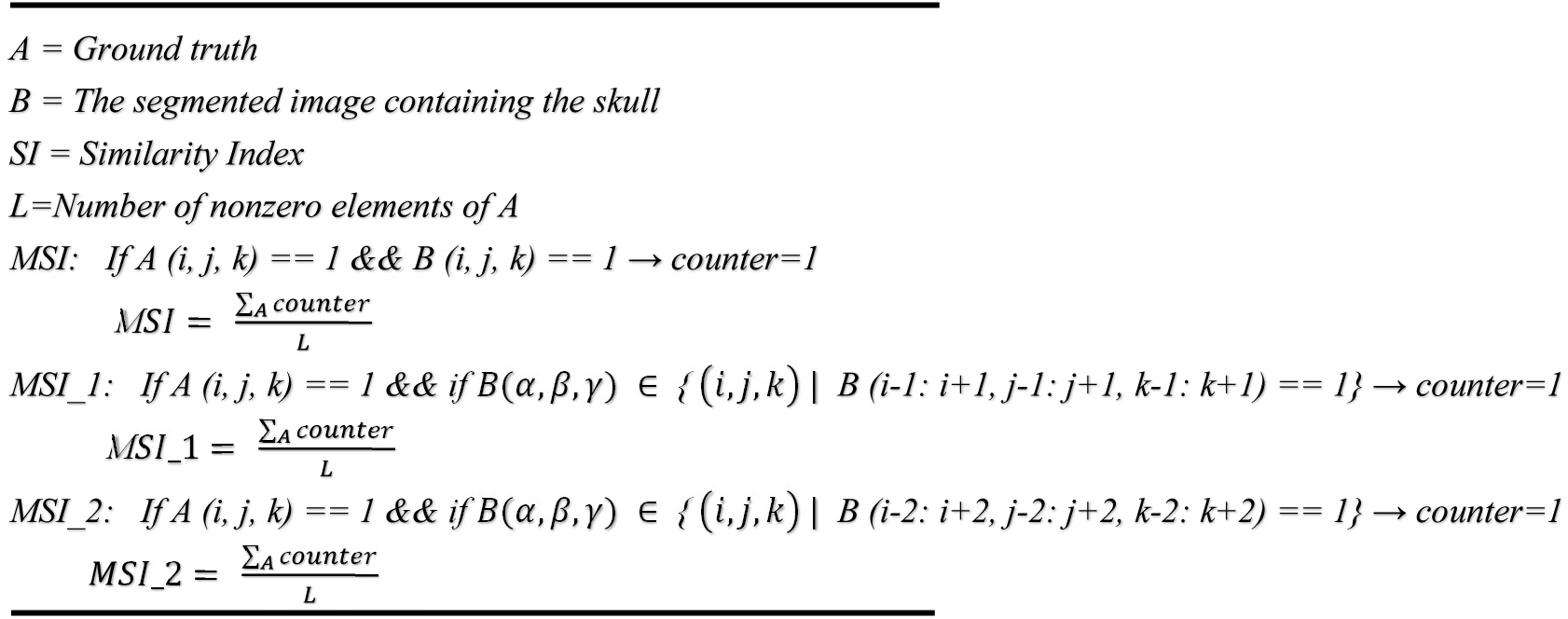

#### 2.4.3 Evaluation of the Results Using an MR-CT Image

The skull segmentation method can also be evaluated using MR and CT images obtained from a neonate in a short period of time. The CT image must be in the same space as the MR image to be evaluated. The multimodal registration method described in Section 2.3.2 is used for this purpose. The cranial bones are then extracted from the CT image using thresholding and used as GT. The DSC and MHD are used for comparison (Dice, 1945; Dubuisson & Jain, 1994). The CT and MR images in this experiment were obtained from a subject at 40and 42 weeks GA, respectively. Figure 11 depicts the test subject’s CT and MR images after registration.

**Fig. 11.**
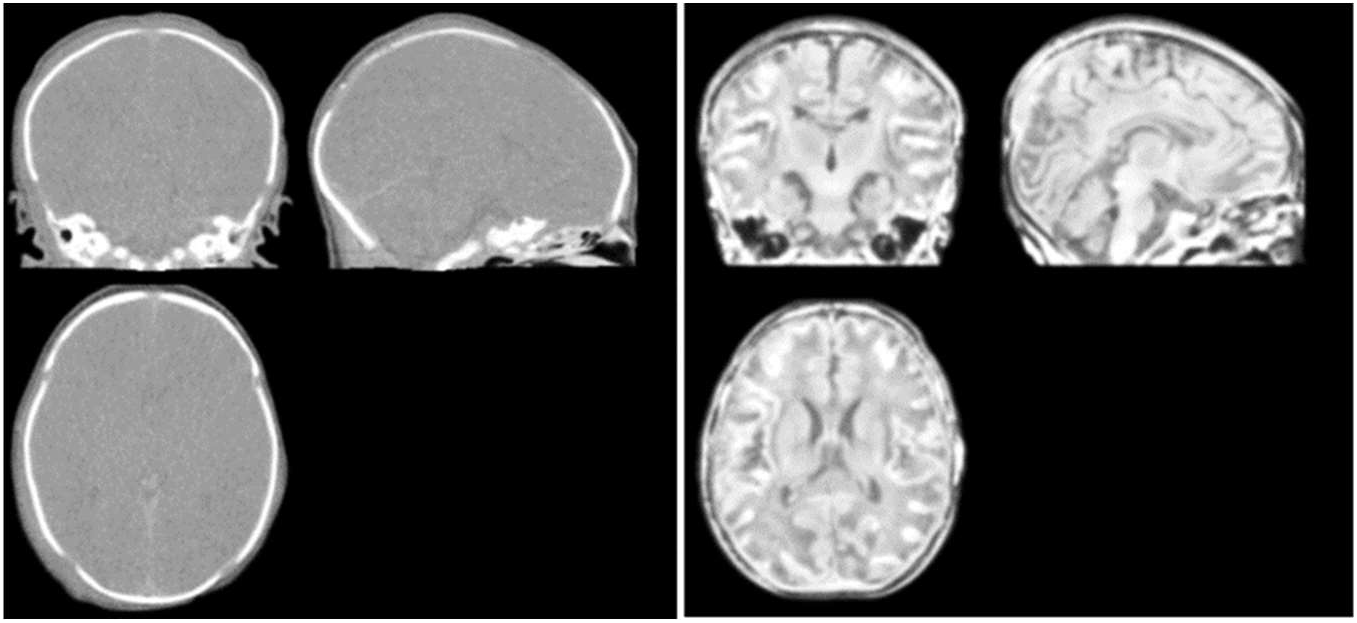
The corresponding CT and MR images of the subject Test1 after multimodal registration

## 3. Results

All experiments were carried out on a personal computer equipped with an Intel Core i7-11800H CPU and an NVIDIA Geforce RTX 3050ti GPU. Skull segmentation from an input MR image takes about 43 minutes. It is the result of several processes: Creation of subject-specific MR atlases (Block I), subject specific CT atlases (Block II), neonatal skull and scalp probability maps (Block III) take 7, 34, and 1 minute(s), respectively. Skull segmentation from the input MR image in FSL takes less than a minute, and intracranial extraction takes less than 30 seconds.

### 3.1 Evaluation of the Results Using Intracranial Tissue

As described in Section 2.4.2, we extracted intracranial tissues for each test image. The proposed method was then compared with the well-known BET (Smith, 2002) and BSE (Shattuck et al., 2001) methods. Our algorithm outperformed other methods in terms of accuracy: The obtained Dice coefficient, modified Hausdorff distance, sensitivity, and specificity were 96.31%, 6.27 mm, 98.23%, and 97.39%, respectively. Our method’s Dice coefficients were significantly higher when compared to all other methods. Figure 12 depicts box plots with various metric values for the evaluated methods on test images. Table 1 expresses the evaluation metrics’ means and standard deviations (SD). Figure 13 shows sample outputs from each method, specifically the case with the high Dice coefficient. We employed the same parameters for BSE (Shattuck et al., 2001) as Serag et al. (Serag et al., 2016). The default parameters were used for BET (Smith, 2002).

**Fig. 12.**
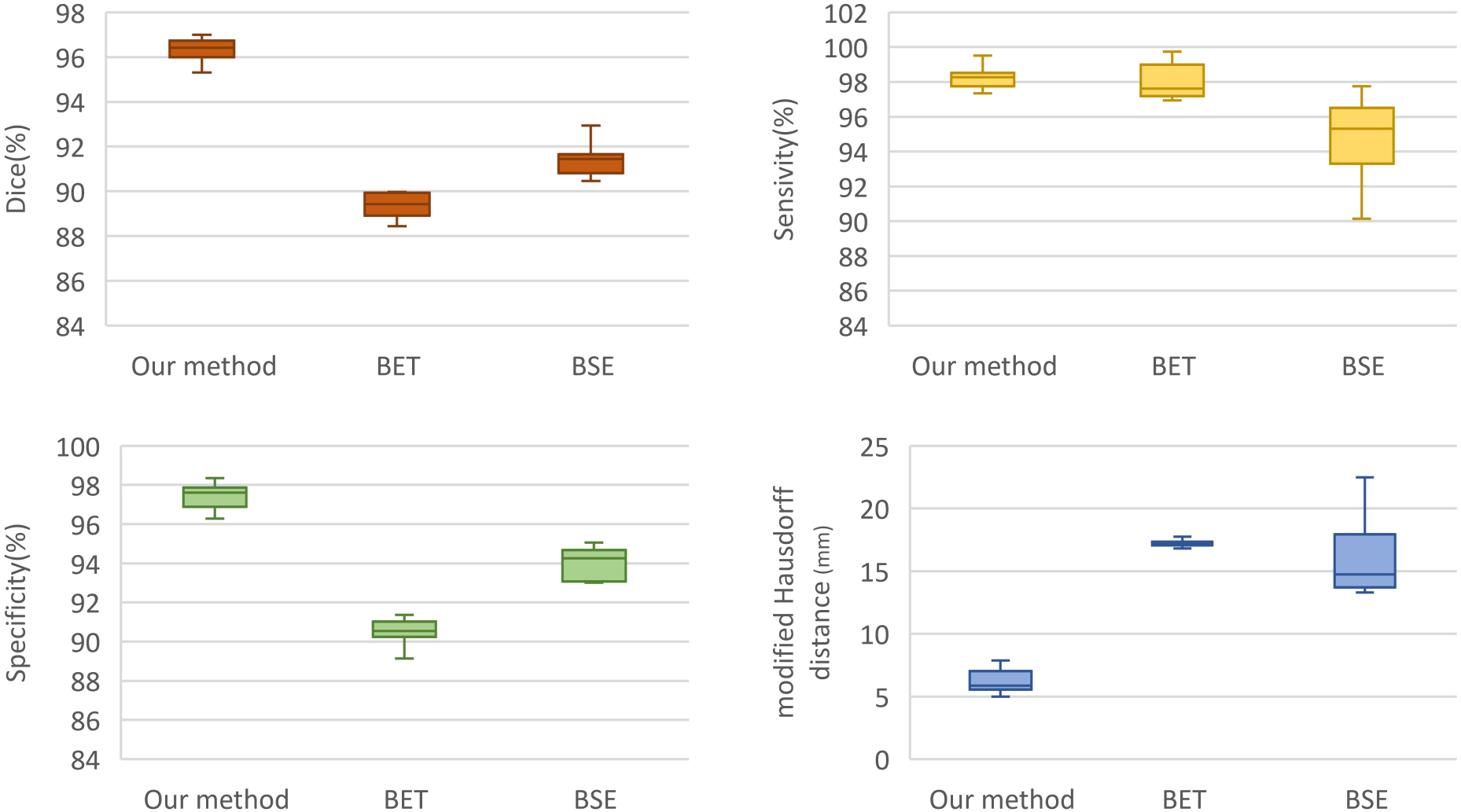
Box plots of Dice coefficient, sensitivity, and specificity and modified Hausdorff distance

**Fig. 13.**
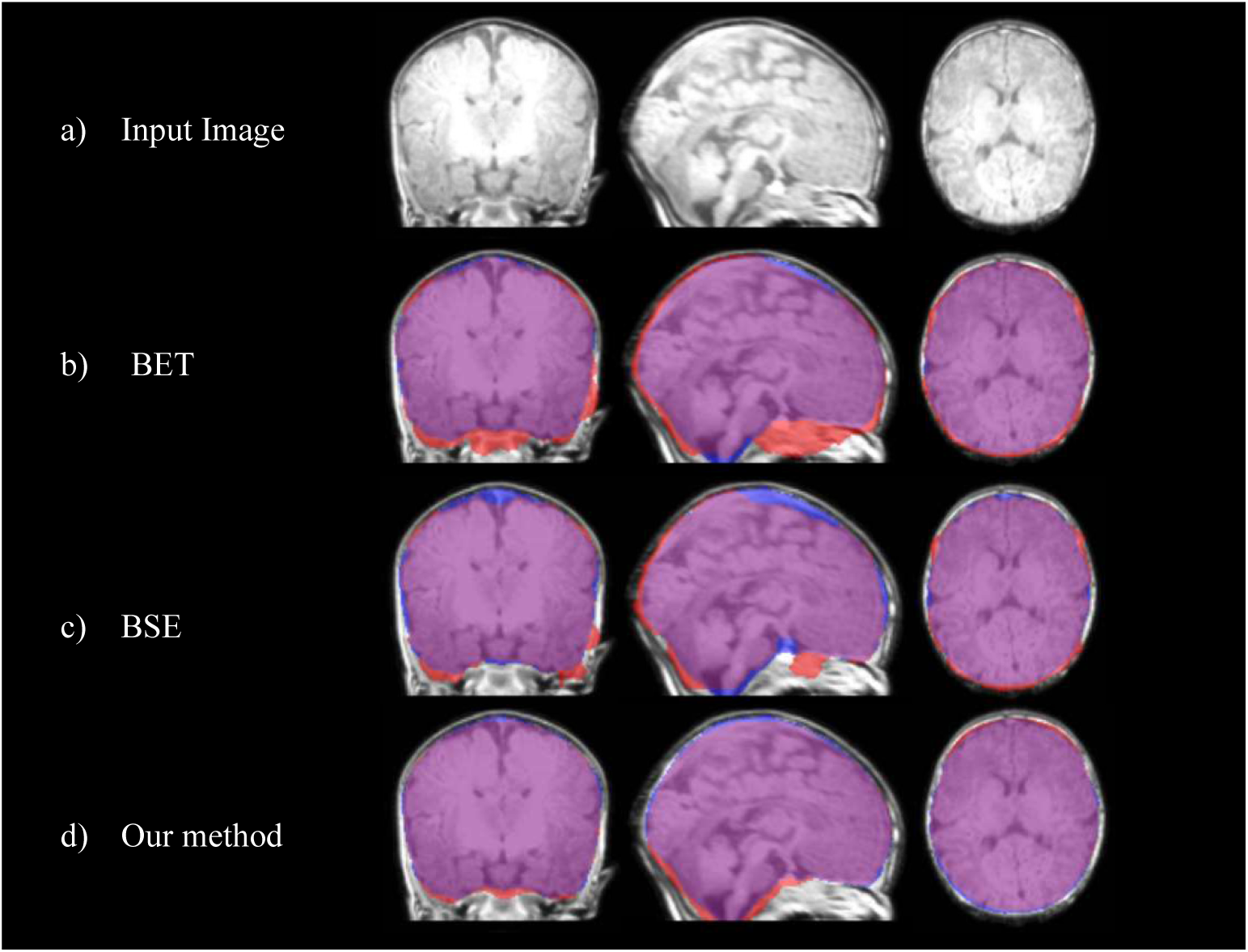
Typical brain extraction results for different methods. For each method, Blue: GT; Red: Automatic segmentation; Violet: Overlap between GT and automatic segmentation. a) MRI of subject N6, b) BET, c) BSE, d) Our method

**Table 1.**
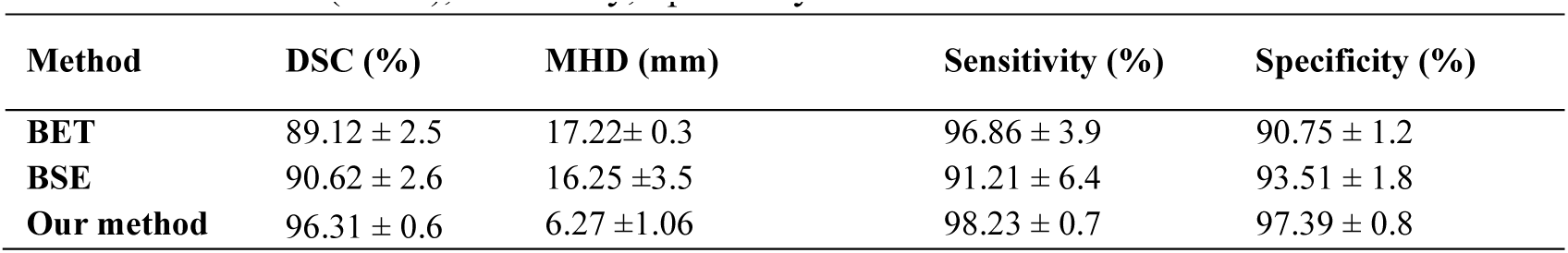
Mean and standard deviation (SD) of the evaluation metrics: Dice coefficient (DSC), modified Hausdorff distance (MHD), Sensitivity, Specificity.

| Method | DSC (%) | MHD (mm) | Sensitivity (%) | Specificity (%) |
| --- | --- | --- | --- | --- |
| **BET** | 89.12 ± 2.5 | 17.22± 0.3 | 96.86 ± 3.9 | 90.75 ± 1.2 |
| **BSE** | 90.62 ± 2.6 | 16.25 ±3.5 | 91.21 ± 6.4 | 93.51 ± 1.8 |
| **Our method** | 96.31 ± 0.6 | 6.27 ±1.06 | 98.23 ± 0.7 | 97.39 ± 0.8 |

### 3.2 Evaluation of the Results Using 3D Anatomical Landmarks

Figure 14 illustrates the alignment of extracted anatomical landmarks (anterior and posterior sutures) and cranial bones segmented by the proposed algorithm. Table 2’s second, fourth, and sixth columns indicate the evaluation of segmented MR images using the MSI, MSI_1, and MSI_2 indices. Obviously, values closer to one represent more accurate results. It is worth noting that these values represent the shape similarity between reconstructed sutures and corresponding landmarks. The low MSI values in Table 2 are primarily due to the proposed algorithm extracting the coronal and lambdoid sutures as closed. Table 2 shows the results of Ghadimi’s algorithm (Ghadimi, 2017) in the third, fifth, and seventh columns. They are generally lower than ours, demonstrating the proposed method’s superiority. Furthermore, the surface of the skull in Ghadimi’s method (Ghadimi, 2017) had many protrusions and depressions, as well as discontinuities, which were improved in this study.

**Fig. 14.**
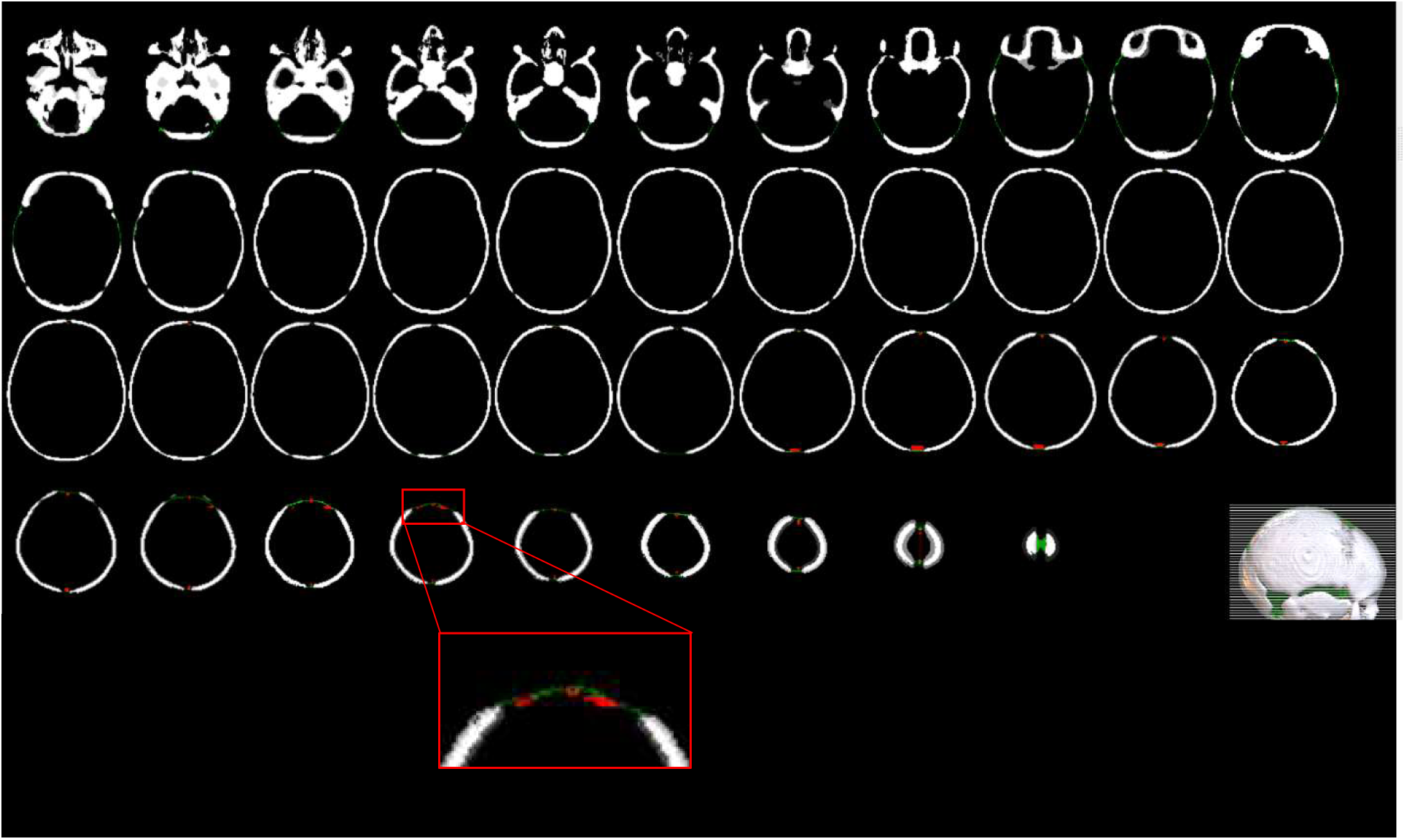
Axial slices of the skull extracted from the N1 subject image. Red tracks in each slice show the anatomical landmarks corresponding to sutures. Green tracks in each slice show the fontanels reconstructed using the coupled level set algorithm (Ghadimi et al., 2015).

**Table 2.** Evaluation of the proposed skull segmentation method using MSI, MSI-1 and MSI-2 similarity coefficients.

| Subjects | MSI | MSI* | MSI-1 | MSI-1* | MSI-2 | MSI-2* |
| --- | --- | --- | --- | --- | --- | --- |
| N1 | 0.23 | 0.05 | 0.85 | 0.28 | 0.97 | 0.33 |
| N2 | 0.13 | 0.06 | 0.56 | 0.30 | 0.78 | 0.41 |
| N3 | 0.10 | 0.06 | 0.61 | 0.23 | 0.87 | 0.31 |
| N4 | 0.03 | 0 | 0.38 | 0.12 | 0.92 | 0.41 |
| N5 | 0.13 | NaN** | 0.41 | NaN** | 0.70 | NaN** |
| N6 | 0.02 | 0 | 0.63 | 0.18 | 0.97 | 0.53 |
| N7 | 0.34 | 0.08 | 0.92 | 0.25 | 0.97 | 0.30 |
| T1 | 0.18 | 0.11 | 0.85 | 0.37 | 0.93 | 0.48 |
| T2 | 0.30 | 0.08 | 0.89 | 0.32 | 0.99 | 0.41 |
| T3 | 0.24 | 0.12 | 0.87 | 0.41 | 0.92 | 0.48 |
| T4 | 0.10 | 0.03 | 0.27 | 0.13 | 0.42 | 0.18 |
| T5 | 0.03 | NaN** | 0.20 | NaN** | 0.71 | NaN** |
| T6 | 0.11 | NaN** | 0.50 | NaN** | 0.88 | NaN** |
| T7 | 0.31 | NaN** | 0.91 | NaN** | 0.97 | NaN** |
| **Average** | **0.16 ± 0.11** | **0.06 ± 0.04** | **0.63 ± 0.25** | **0.26 ± 0.1** | **0.86 ± 0.16** | **0.38 ± .11** |

\* Result obtained by the method in (Ghadimi, 2017)
\*\* No information available

### 3.3 Evaluation Using an MR-CT Image

Figure 15 depicts the superposition of a skull extracted from an MR image (white) and one extracted from a CT image (red). Probabilities less than 0.5 were treated as zero in the probability models developed for segmenting this MR image. DSC is 0.50 and MHD is 1.60 mm between skulls extracted from MR and CT images. This subject’s CT scan was performed two weeks before the MR scan. As a result, based on the World Health Organization’s growth charts of neonatal head circumference from birth to 13 weeks (Mercedes de Onis et al, 2007), we expect the head circumference to be larger in the MR image due to the infant’s growth (Figure 15). Therefore, the difference in skull size between CT and MR images helps to explain the low DSC value. In the CT and MR images, the head circumference is 355 mm and 368 mm, respectively. According to the WHO Child Growth Standards (Mercedes de Onis et al, 2007), the average head circumference difference during this period is around 1.5 cm, so the difference of 1.3 cm appears acceptable.

**Fig. 15.**
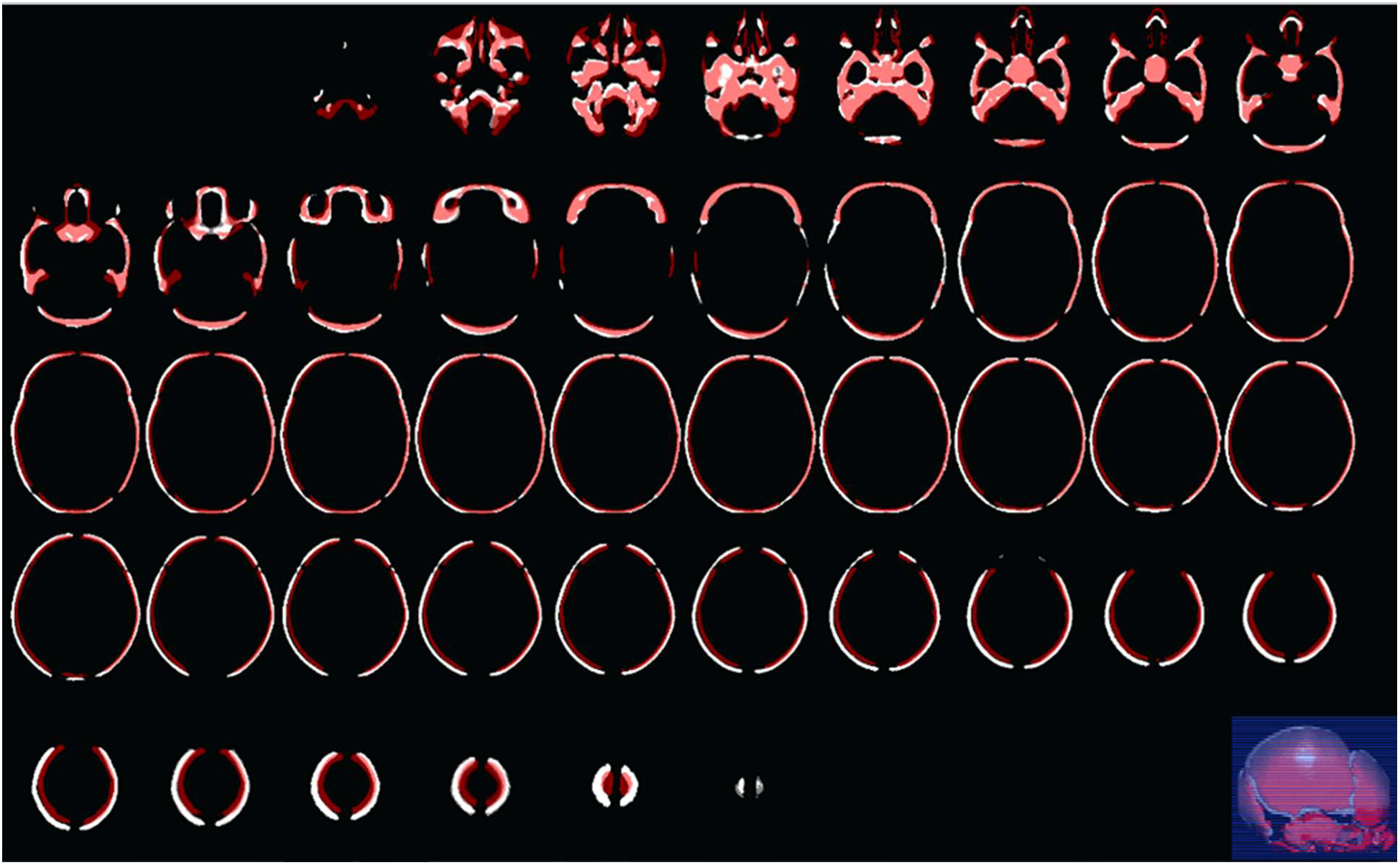
Evaluation of the skull segmentation method using CT and MR data acquired from a neonate in 40- and 42-weeks GA, respectively. The extracted skull from MR image (white) is superposed on the extracted skull from corresponding CT image (red).

## 4. Discussion

The effect of fontanels and sutures on EEG and MEG source analysis using realistic neonatal head models was investigated by several studies (Azizollahi et al., 2016; Dehaes et al., 2013; Lew et al., 2013; Noreika et al., 2020; Odabaee et al., 2014; Roche-Labarbe et al., 2008). Realistic models of a neonate’s head created by MR imaging are commonly used to identify different tissues in the head that exhibit different electrical and optical properties. Accordingly, accurate skull segmentation of neonatal cerebral MRIs can have a significant impact on the results of neonatal functional analysis using EEG, MEG, near infrared spectroscopy (NIRS) and functional MRI (fMRI). Unfortunately, the MRI data does not provide perceivable information needed to localize the cranial bones in the neonatal head image. Accordingly, a bimodal approach based on MRI and CT images of infants’ heads was proposed for the first time in this paper to solve the challenging problem of skull segmentation from MRIs. The proposed approach relies on retrospective MR and CT images to create subject-specific atlases of the neonatal head, including probability maps of the skull, scalp, and CSF. These subject-specific atlases are employed by an EM algorithm in conjunction with a Markov random field model to segment the skull from neonatal head MRI data acquired from a subject. Our approach consisted of three main blocks. In the first block, subject specific head and intracranial templates and CSF probability maps are created using retrospective MR data acquired from neonates of 39-42 weeks GA. In the second block, the head and intracranial MR templates are employed to create a subject-specific CT head template using retrospective CT data acquired from neonates in the same age range. The third block uses the CT head template to create subject-specific probability maps of the skull and scalp. Finally, the subject-specific CSF, skull, and scalp probability maps are fed to an EM algorithm in conjunction with Markov random field model, as implemented in FSL software, to segment the scalp, skull, and intracranial from the input MR image. The proposed approach provided realistic segmentation of the skull in which fontanels and sutures were taken into consideration.

The evaluation of the proposed neonatal skull segmentation approach was another challenge to overcome due to the lack of GT. Former methods for neonatal skull segmentation from MRI have often used manual or semi-automatic segmentation of image slices as GT (Mahapatra, 2012, Shi et al., 2012, Yamaguchi et al., 2010). In some studies, only a few slices have been segmented and used as GT due to the invisibility of cranial bones in certain slices and the time-consuming process of manual segmentation. On the other hand, most of these methods, ignored sutures and fontanels (Despotovic et al., 2009, Burguet et al., 2004). In other words, they considered the skull as a continuous and homogeneous tissue which was far from a realistic skull geometry in neonates (Despotovic et al., 2009, Burguet et al., 2004). The issues underscored the need for a trusted GT to assess the results. We noticed that despite the quasi-invisibility of cranial bones, the sutures are perceivable as bright regions in the three-dimensional volume rendering of the neonatal head in T1-weighted MR images. Therefore, they were considered as three-dimensional landmarks and used as GT. Accordingly, we reconstructed the sutures from skull segmentation results and then compared them with the sutures which had been extracted manually from the original MR data as GT. For this purpose, modified versions of the Dice similarity coefficient were introduced throughout this paper.

Another method for evaluating the results is to extract intracranial tissues from MRIs and compare them to GT. Comparisons demonstrated that our algorithm outperforms well-known skull stripping methods in terms of different criteria, including the Dice index, modified Hausdorff distance, sensitivity, and specificity. This outperformance is mainly due to our bimodal MR-CT atlas-based segmentation approach.

An alternative reliable solution to evaluate our neonatal skull segmentation method is appealing to retrospective bimodal MR-CT head data from the same neonates that have been acquired within a short time interval. Obviously, access to such bimodal data is very restricted due to several factors. First, acquiring CT images from neonates is rarely authorized when there are strong clinical reasons. Second, finding bimodal MR-CT data that have been acquired from neonates with normal head anatomy is challenging because in most cases, the subjects were suffering from some kind of abnormality in their head such as bleeding, concussion, etc. Third, the time interval between MR and CT acquisition must be sufficiently short due to the rapid development of the skull in neonates. In this paper, the bimodal data acquired from only one subject were employed for evaluating the proposed neonatal skull segmentation method. Taking advantage of more bimodal test data would be helpful for an exhaustive evaluation of the proposed method.

The anterior fontanel size is defined as the average of its small and large diameters. The normal size of the anterior fontanel in infants at birth is between 0.6 and 3.6 cm (Kiesler & Ricer, 2003). In this study, the average value of the anterior fontanel size on our MR dataset was 3.19 cm, which falls within the normal range (Mohtasebi et al., 2021). The skull thickness is another criterion that can be used to validate the proposed method. Li et al. (Li et al., 2015) demonstrated that the thickness of the skull gradually and inhomogeneously increases from birth. The average thickness of extracted bones from 14 newborn MR images was 1.88 mm, which is close to the 2.16 mm reported for healthy infants by (Li et al., 2015).

The proposed method outperformed its close counterparts in terms of modified similarity indices, notably MSI_2. The creation and use of subject-specific atlases justify the superiority of the proposed approach over Ghadimi’s method (Ghadimi, 2017), which suggests creating atlases in the newborn stereotaxic space (Kazemi et al., 2007). On the other hand, in the method proposed by Kazemi et al. (Kazemi et al., 2008), the skull probability map was created as a continuous structure without considering sutures and fontanels. Obviously, including sutures and fontanels is mandatory to achieve a realistic segmentation of the skull from neonatal cerebral MR images.

In this study, a limited number of retrospective MR and CT images were used to create subject-specific atlases. Obviously, using larger datasets of each modality, which include skulls with diverse morphologies may improve the generalization property of the created atlases. Furthermore, one may use a multi-atlas approach to create a specific atlas that is closer to the subject’s head geometry. However, access to such a large dataset with normal head geometry particularly in CT modality, is a challenging issue. The multimodal atlas data should also include manually extracted tissues such as intracranial, CSF, skull, and scalp for use in the creation of subject-specific atlases and probability maps.

## 5. Conclusion

An atlas-based bimodal MR-CT approach for realistic skull segmentation in neonatal MR data was proposed in this study. The new approach deploys preprocessed retrospective MR and CT head data to create subject-specific MR and CT head templates. It also creates subject-specific CSF, skull, and scalp probability maps using semiautomatically extracted CSF (from atlas MRIs), skull, and scalp (from atlas CT scans) tissues. The resulting subject-specific MR atlas and CSF, skull, and scalp probability maps are used within the framework of the EM algorithm in conjunction with a Markov random field model to segment the skull from an input neonatal head MRI. The proposed approach provided realistic skull segmentation results that included sutures and fontanels. The evaluation of the results using three elaborate evaluation procedures demonstrated the good performance of the proposed algorithm. The proposed skull segmentation method seems promising in neurodevelopmental and neuroscience studies.

## Acknowledgments

We would like to thank Dr. Sona Ghadimi for her comments on the initial phase of research. We would particularly like to thank the Cognitive Sciences and Technologies Council (COGC), Iran, in the framework of the Neurobiom project number 96P97, for their support and guidance.

## References

1. Antonakakis M, et al. The effect of stimulation type, head modeling, and combined EEG and MEG on the source reconstruction of the somatosensory P20/N20 component. Hum Brain Mapp 2019;40(17):5011–28. http://dx.doi.org/10.1002/hbm.24754

2. Avants, B. B., Epstein, C. L., Grossman, M., & Gee, J. C. (2008). Symmetric diffeomorphic image registration with cross-correlation: evaluating automated labeling of elderly and neurodegenerative brain. Med Image Anal, 12(1), 26–41. https://doi.org/10.1016/j.media.2007.06.004

3. Avants, B. B., Tustison, N. J., Stauffer, M., Song, G., Wu, B., & Gee, J. C. (2014). The Insight ToolKit image registration framework. Front Neuroinform, 8, 44. https://doi.org/10.3389/fninf.2014.00044

4. Azizollahi, H., Aarabi, A., & Wallois, F. (2016). Effects of uncertainty in head tissue conductivity and complexity on EEG forward modeling in neonates. Hum Brain Mapp, 37(10), 3604–3622. https://doi.org/10.1002/hbm.23263

5. Burguet, J., Gadi, N., & Bloch, I. (2004). Realistic models of children heads from 3D-MRI segmentation and tetrahedral mesh construction. Proceedings. 2nd International Symposium on 3D Data Processing, Visualization and Transmission, 2004. 3DPVT 2004., https://doi.org/10.1109/TDPVT.2004.1335298

6. Cardoso, M. J., Melbourne, A., Kendall, G. S., Modat, M., Robertson, N. J., Marlow, N., & Ourselin, S. (2013). AdaPT: an adaptive preterm segmentation algorithm for neonatal brain MRI. NeuroImage, 65, 97–108. https://doi.org/10.1016/j.neuroimage.2012.08.009

7. Cherel, M., Budin, F., Prastawa, M., Gerig, G., Lee, K., Buss, C., Lyall, A., Consing, K. Z., & Styner, M. (2015). Automatic tissue segmentation of neonate brain MR Images with subject-specific atlases. Medical Imaging 2015: Image Processing, https://doi.org/10.1117/12.2082209

8. Daliri, M., Moghaddam, H. A., Ghadimi, S., Momeni, M., Harirchi, F., & Giti, M. (2010). Skull segmentation in 3D neonatal MRI using hybrid Hopfield Neural Network. 2010 Annual International Conference of the IEEE Engineering in Medicine and Biology, https://doi.org/10.1109/iembs.2010.5627619

9. Dehaes, M., Kazemi, K., Pelegrini-Issac, M., Grebe, R., Benali, H., & Wallois, F. (2013). Quantitative effect of the neonatal fontanel on synthetic near infrared spectroscopy measurements. Hum Brain Mapp, 34(4), 878–889. https://doi.org/10.1002/hbm.21483

10. Despotovic, I., Deburchgraeve, W., Hallez, H., Vansteenkiste, E., & Philips, W. (2009). Development of a realistic head model for EEG event-detection and source localization in newborn infants. 2009 Annual International Conference of the IEEE Engineering in Medicine and Biology Society, https://doi.org/10.1109/IEMBS.2009.5335052

11. Dice, L. R. (1945). Measures of the amount of ecologic association between species. Ecology, 26(3), 297–302. https://doi.org/10.2307/1932409

12. Dubuisson, M.-P., & Jain, A. K. (1994). A modified Hausdorff distance for object matching. Proceedings of 12th international conference on pattern recognition, https://doi.org/10.1109/ICPR.1994.576361

13. Fedorov, A., Beichel, R., Kalpathy-Cramer, J., Finet, J., Fillion-Robin, J.-C., Pujol, S., Bauer, C., Jennings, D., Fennessy, F., & Sonka, M. (2012). 3D Slicer as an image computing platform for the Quantitative Imaging Network. Magnetic resonance imaging, 30(9), 1323–1341. https://doi.org/10.1016/j.mri.2012.05.001

14. Gao, Y., Li, J., Xu, H., Wang, M., Liu, C., Cheng, Y., … & Li, X. (2019). A multi-view pyramid network for skull stripping on neonatal T1-weighted MRI. Magnetic resonance imaging, 63, 70–79. https://doi.org/10.1016/j.mri.2019.08.025

15. Ghadimi, S. (2017). Creation of 3D neonatal skull model using MR and CT images. K. N. Toosi University of technology and university of Picardie Jules Verne, Doctoral dissertation.

16. Ghadimi, S., Moghaddam, H. A., Grebe, R., & Wallois, F. (2015). Skull segmentation and reconstruction from newborn CT images using coupled level sets. IEEE journal of biomedical and health informatics, 20(2), 563–573. https://doi.org/10.1109/JBHI.2015.2391991

17. Ghadimi, S., Mohtasebi, M., Abrishami Moghaddam, H., Grebe, R., Gity, M., & Wallois, F. (2017). A Neonatal Bimodal MR-CT Head Template. PloS one, 12(1), e0166112. https://doi.org/10.1371/journal.pone.0166112

18. Gilmore JH, Knickmeyer RC, Gao W. Imaging structural and functional brain development in early childhood. Nat Rev Neurosci 2018;19(3):123–37. https://doi.org/10.1038/nrn.2018.1

19. Gousias, I. S., Edwards, A. D., Rutherford, M. A., Counsell, S. J., Hajnal, J. V., Rueckert, D., & Hammers, A. (2012). Magnetic resonance imaging of the newborn brain: manual segmentation of labelled atlases in term-born and preterm infants. Neuroimage, 62(3), 1499–1509. https://doi.org/10.1016/j.neuroimage.2012.05.083

20. Henkelman, R. M., Watts, J. F., & Kucharczyk, W. (1991). High signal intensity in MR images of calcified brain tissue. Radiology, 179(1), 199–206. https://doi.org/10.1148/radiology.179.1.1848714

21. Kazemi, K., Ghadimi, S., Abrishami-Moghaddam, H., Grebe, R., Gondry-Jouet, C., & Wallois, F. (2008). Neonatal probabilistic models for brain, CSF and skull using T1-MRI data: preliminary results. 2008 30th Annual International Conference of the IEEE Engineering in Medicine and Biology Society, https://doi.org/10.1109/IEMBS.2008.4650060

22. Kazemi, K., Moghaddam, H. A., Grebe, R., Gondry-Jouet, C., & Wallois, F. (2007). A neonatal atlas template for spatial normalization of whole-brain magnetic resonance images of newborns: preliminary results. Neuroimage, 37(2), 463–473. https://doi.org/10.1016/j.neuroimage.2007.05.004

23. Keith A. Johnson, M.D. Brigham and Womenśs Hospital, Harvard Medical School https://www.med.harvard.edu/aanlib/hem.html. (accessed 23 April 2022)

24. Kiesler, J., & Ricer, R. (2003). The abnormal fontanel. Am Fam Physician, 67(12), 2547–2552. https://www.ncbi.nlm.nih.gov/pubmed/12825844

25. Kobashi, S., & Udupa, J. K. (2013, July). Fuzzy connectedness image segmentation for newborn brain extraction in MR images. In 2013 35th Annual international conference of the IEEE engineering in medicine and biology society (EMBC) (pp. 7136–7139). IEEE. https://doi.org/10.1109/embc.2013.6611203

26. Lemieux, L., Wieshmann, U. C., Moran, N. F., Fish, D. R., & Shorvon, S. D. (1998). The detection and significance of subtle changes in mixed-signal brain lesions by serial MRI scan matching and spatial normalization. Med Image Anal, 2(3), 227–242. https://doi.org/10.1016/s1361-8415(98)80021-2

27. Lew, S., Sliva, D. D., Choe, M. S., Grant, P. E., Okada, Y., Wolters, C. H., & Hamalainen, M. S. (2013). Effects of sutures and fontanels on MEG and EEG source analysis in a realistic infant head model. Neuroimage, 76, 282–293. https://doi.org/10.1016/j.neuroimage.2013.03.017

28. Li, Z., Park, B. K., Liu, W., Zhang, J., Reed, M. P., Rupp, J. D., Hoff, C. N., & Hu, J. (2015). A statistical skull geometry model for children 0-3 years old. PloS one, 10(5), e0127322. https://doi.org/10.1371/journal.pone.0127322

29. Li G, Wang L, Yap PT, Wang F, Wu Z, Meng Y, et al. Computational neuroanatomy of baby brains: a review. Neuroimage 2019;185(1):906–25. https://doi.org/10.1016/j.neuroimage.2018.03.042

30. Mahapatra, D. (2012). Skull stripping of neonatal brain MRI: using prior shape information with graph cuts. Journal of digital imaging, 25(6), 802–814. https://doi.org/10.1007/s10278-012-9460-z

31. Makropoulos, A., Counsell, S. J., & Rueckert, D. (2018). A review on automatic fetal and neonatal brain MRI segmentation. NeuroImage, 170, 231–248. https://doi.org/10.1016/j.neuroimage.2017.06.074

32. Mohtasebi, M., Bayat, M., Ghadimi, S., Moghaddam, H. A., & Wallois, F. (2021). Modeling of Neonatal Skull Development Using Computed Tomography Images. Irbm, 42(1), 19–27. https://doi.org/10.1016/j.irbm.2020.02.002

33. Nielsen, J. D., Madsen, K. H., Puonti, O., Siebner, H. R., Bauer, C., Madsen, C. G., Saturnino, G. B., & Thielscher, A. (2018). Automatic skull segmentation from MR images for realistic volume conductor models of the head: Assessment of the state-of-the-art. Neuroimage, 174, 587–598. https://doi.org/10.1016/j.neuroimage.2018.03.001

34. Noorizadeh, N., Kazemi, K., Danyali, H., & Aarabi, A. (2019). Multi-atlas based neonatal brain extraction using a two-level patch-based label fusion strategy. Biomedical Signal Processing and Control, 54, 101602. https://doi.org/10.1016/j.bspc.2019.101602

35. Noreika, V., Georgieva, S., Wass, S., & Leong, V. (2020). 14 challenges and their solutions for conducting social neuroscience and longitudinal EEG research with infants. Infant Behavior and Development, 58, 101393. https://doi.org/10.1016/j.infbeh.2019.101393

36. Odabaee, M., Tokariev, A., Layeghy, S., Mesbah, M., Colditz, P. B., Ramon, C., & Vanhatalo, S. (2014). Neonatal EEG at scalp is focal and implies high skull conductivity in realistic neonatal head models. Neuroimage, 96, 73–80. https://doi.org/10.1016/j.neuroimage.2014.04.007

37. Okada, E., & Delpy, D. T. (2003). Near-infrared light propagation in an adult head model. I. Modeling of low-level scattering in the cerebrospinal fluid layer. Appl Opt, 42(16), 2906–2914. https://doi.org/10.1364/ao.42.002906

38. Otsu, N. (1979). A threshold selection method from gray-level histograms. IEEE transactions on systems, man, and cybernetics, 9(1), 62–66. https://doi.org/10.1109/TSMC.1979.4310076

39. Péeportée, M., Ilea Ghita, D. E., Twomey, E., and Whelan, P. F. (2011). A Hybrid Approach to Brain Extraction from Premature Infant MRI. In Image Analysis, volume 6688, pages 719–730. Springer Berlin Heidelberg, Berlin, Heidelberg. http://dx.doi.org/10.1007/978-3-642-21227-7_67

40. Prastawa, M., Gilmore, J. H., Lin, W., & Gerig, G. (2005). Automatic segmentation of MR images of the developing newborn brain. Med Image Anal, 9(5), 457–466. https://doi.org/10.1016/j.media.2005.05.007

41. Richter, L., & Fetit, A. E. (2022). Accurate segmentation of neonatal brain MRI with deep learning. Frontiers in Neuroinformatics, 16. https://doi.org/10.3389/fninf.2022.1006532

42. Roche-Labarbe, N., Aarabi, A., Kongolo, G., Gondry-Jouet, C., Dumpelmann, M., Grebe, R., & Wallois, F. (2008). High-resolution electroencephalography and source localization in neonates. Hum Brain Mapp, 29(2), 167–176. https://doi.org/10.1002/hbm.20376

43. Rorden, C., Bonilha, L., Fridriksson, J., Bender, B., & Karnath, H. O. (2012). Age-specific CT and MRI templates for spatial normalization. Neuroimage, 61(4), 957–965. https://doi.org/10.1016/j.neuroimage.2012.03.020

44. Serag, A., Blesa, M., Moore, E. J., Pataky, R., Sparrow, S. A., Wilkinson, A. G., … & Boardman, J. P. (2016). Accurate Learning with Few Atlases (ALFA): an algorithm for MRI neonatal brain extraction and comparison with 11 publicly available methods. Scientific Reports, 6(1), 1–15. https://doi.org/10.1038/srep23470

45. Shattuck, D. W., Sandor-Leahy, S. R., Schaper, K. A., Rottenberg, D. A., & Leahy, R. M. (2001). Magnetic resonance image tissue classification using a partial volume model. NeuroImage, 13(5), 856–876. https://doi.org/10.1006/nimg.2000.0730

46. Shi, F., Wang, L., Dai, Y., Gilmore, J. H., Lin, W., & Shen, D. (2012). LABEL: pediatric brain extraction using learning-based meta-algorithm. Neuroimage, 62(3), 1975–1986. https://doi.org/10.1016/j.neuroimage.2012.05.042

47. Smith, S. M. (2002). Fast robust automated brain extraction. Human brain mapping, 17(3), 143–155. https://doi.org/10.1002/hbm.10062

48. Smith, S. M., Jenkinson, M., Woolrich, M. W., Beckmann, C. F., Behrens, T. E., Johansen-Berg, H., Bannister, P. R., De Luca, M., Drobnjak, I., & Flitney, D. E. (2004). Advances in functional and structural MR image analysis and implementation as FSL. Neuroimage, 23, S208–S219. https://doi.org/10.1016/j.neuroimage.2004.07.051

49. Tustison, N. J., Avants, B. B., Cook, P. A., Zheng, Y., Egan, A., Yushkevich, P. A., & Gee, J. C. (2010). N4ITK: improved N3 bias correction. IEEE Trans Med Imaging, 29(6), 1310–1320. https://doi.org/10.1109/TMI.2010.2046908

50. Wang, G., Hu, Y., Li, X., Wang, M., Liu, C., Yang, J., & Jin, C. (2020). Impacts of skull stripping on construction of three-dimensional T1-weighted imaging-based brain structural network in full-term neonates. Biomedical engineering online, 19(1), 1–13. https://doi.org/10.1186/s12938-020-00785-0

51. Weinreb, J. C., Brateman, L., Babcock, E. E., Maravilla, K. R., Cohen, J. M., & Horner, S. (1985). Chemical shift artifact in clinical magnetic resonance images at 0.35 T. American journal of roentgenology, 145(1), 183–185. https://doi.org/10.2214/ajr.145.1.183

52. Mercedes de Onis et al. (2007). WHO child growth standards: head circumference-for-age, arm circumference-for-age, triceps skinfold-for-age and subscapular skinfold-for-age: methods and development. ISBN 978 92 4 154718 5. Downloadable from https://www.who.int/publications/i/item/9789241547185. (accessed 23 April 2022)

53. Yamaguchi, K., Fujimoto, Y., Kobashi, S., Wakata, Y., Ishikura, R., Kuramoto, K., Imawaki, S., Hirota, S., & Hata, Y. (2010). Automated fuzzy logic-based skull stripping in neonatal and infantile MR images. International Conference on Fuzzy Systems, https://doi.org/10.1109/FUZZY.2010.5584839

